# De-novo design of actively spinning and gyrating spherical micro-vesicles

**DOI:** 10.1101/2024.09.17.613442

**Authors:** Veerpal Kaur, Subhashree S. Khuntia, Charu Taneja, Abhishek Chaudhuri, K. P. Yogendran, Sabyasachi Rakshit

**Affiliations:** Department of Chemical Sciences, Indian Institute of Science Education and Research Mohali,140306, Punjab, India; Department of Physical Sciences, Indian Institute of Science Education and Research Mohali, Punjab, India

## Abstract

Engineering spherical self-propelled swimmers that exhibit rotation and directed translation has posed a significant experimental challenge in biomedicine design. Often a secondary external field or asymmetric geometry is employed to generate rotation, complicating the design process. In this work, we utilize spherical Giant Unilamellar Vesicles (GUVs) as chassis and enzymes undergoing cyclic, non-reciprocal conformational changes as power units to establish design principles to synthesize autonomous spherical micro(µ)-rotors. Leveraging transient interactions, we induce spontaneous symmetry-breaking in enzyme distribution on GUVs, enabling diverse movements from pure spinning to spiral 3D trajectories. With this design, we now open new avenues for advancing self-propelled systems with biocompatible materials, unlocking innovations in biomedical applications.

## Introduction

Living systems harness energy from their surroundings to achieve complex movements, such as simultaneous rotation and translation(*1*). This capability is particularly evident in microorganisms like Chlamydomonas(*2*) and Volvox(*3*, *4*), which navigate challenging environments with remarkable precision. In contrast, synthetic systems have struggled to replicate these feats. The challenge lies in designing self-rotating spherical particles that can autonomously move without relying on external fields(*5–7*) or asymmetric geometries(*8–10*). In this study, we identify design principles for synthesizing autonomous μ-rotors using biocompatible GUVs. Our μ-rotors are spherical and exhibit a variety of swimming dynamics, from pure rotation to complex combinations of spin and translatory motions which persist for hours and range over length scales of millimeters. The spherical shape of GUVs emerges as the preferred geometry in colloids synthesized through solution-based methods. Our research thus has the potential to unlock new applications in synthetic biology(*11–13*), drug delivery(*14*, *15*), and the development of advanced materials that mimic biological systems(*16*). Active spherical rotors are also known to promote collective behavior akin to the dynamic patterns observed in living crystals(*17*), where particle interactions lead to coordinated movement and complex structures(*18*).

These μ-rotors are powered by flexible enzymes embedded on the membrane of GUVs. These enzymes have been shown to exhibit remarkable capabilities in generating motion through catalytic turnover(*19*) when anchored onto solid microscopic beads(*20*, *21*) or nanoscopic vesicles(*22*). However, all such motions observed have been linear. We suggest that a key limitation in these systems was the uncontrolled distribution of enzymes, as well as their arbitrary orientation on spherical surfaces. This randomness prevents the generation of net torque, leading to a lack of spin or rotational movement. To induce autonomous *rotations*, we hypothesize that the condensation of the advection flow generated by flexible enzymes is essential for creating an effective tangential surface velocity field. This is analogous to the coordinated beating of flagella/cilia that generates rotational flow around microorganisms like Volvox(*3*, *4*), Chlamydomonas(*2*), starfish embryo(*17*) etc., causing them to spin as they swim(*23*, *24*).

We propose that this may be achieved by ensuring that the *power sources* are *clustered* on the surface and that the carrier surface is made sufficiently *rigid*. We will now describe our experiments that led us to these inferences.

## Results

### *Flexible* enzymes as force-generators

We selected Sortase A (SrtA) as our model enzyme, using Adenylate Kinase (ADK) and Tobacco Etch Virus protease (TEV) as controls. The rationale for selecting SrtA is twofold. First, SrtA is a flexible enzyme that undergoes simultaneous structural and conformational changes during transpeptidation(*25–27*). Specifically, SrtA transitions from a random coil to 3_10_ helices while also adopting a conformation conducive to peptide bond formation in the presence of a signal peptide (-N term -LPETGSSC-C term-) and a nucleophile (the primary amine of the -GGGGC-chain, polyG) **(Fig. 1a)**. Secondly, SrtA has an intrinsic disordered region (IDR) within the loop between the β6-β7 strands(*28*, *29*) (**Fig. 1a**). IDRs typically facilitate transient protein-protein interactions leading to dynamic clustering(*30*). We utilize this property to generate asymmetrically distributed reaction-clusters on GUVs.

**Fig. 1:**
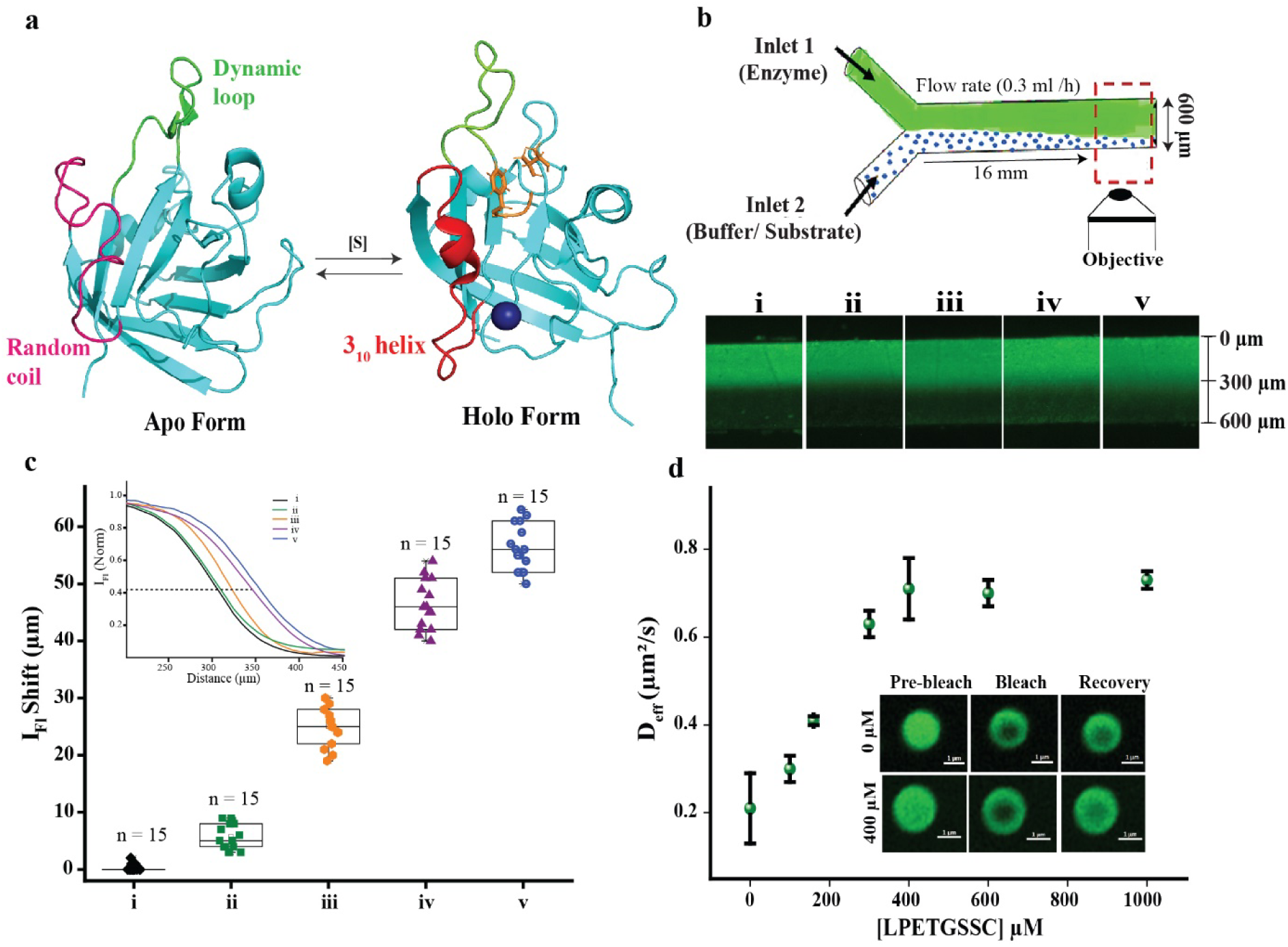
Properties of SrtA. **(a)** Ribbon representation of SrtA in the apo form (PDB: 1IJA) and holo form (PDB: 2KID). In the apo form, dynamic and disordered loops are colored green and magenta, respectively. In the holo form, the red color represents the 3_10_ helix that forms upon binding of the signal peptide (-LPETGSSC-), represented in orange-color. **(b)** The upper panel shows a schematic of a microfluidic setup consisting of two input channels that converge into a single output channel. The lower panel shows the fluorescence images captured 16 mm downstream from the point where the inlets merged, for the following experimental runs: (i) FITC-tagged SrtA at inlet 1 and buffer with only 0.36 mM polyG at inlet 2; (ii) TEV at inlet 1 and substrate (-ENLYFQG-) at inlet 2; (iii) ADK at inlet 1 and substrates (ATP + AMP) at inlet 2; (iv) SrtA at inlet 1 and substrates (0.4 mM of -LPETGSSC-+ 0.36 mM of polyG) at inlet 2; (v) A mixture of SrtA + 0.16 mM of -LPETGSSC-+ 0.36 mM of polyG at inlet 1 and substrates (0.4 mM of -LPETGSSC-+ 0.36 mM of polyG) at inlet 2. **(c)** Fluorescence intensity profiles across the width of the channel output for the experiment runs (i-v) as described in (b) Inset shows the quantitative representation of data for experiments run (i-v) **(d)** Diffusion coefficients of SrtA in coacervates as estimated from FRAP experiments at varying concentrations of the - LPETGSSC-peptide are plotted. Error bars represent statistical variance. (Inset) Representative FRAP images of SrtA in coacervates are shown at two extreme concentrations of -LPETGSSC-: 0 μM (top) and 400 μM (bottom).

ADK also undergoes conformational changes between “closed” and “open” states during reversible phosphate transfer and so may be termed flexible. However, ADK does not possess any IDR(*31*, *32*) (**Fig. S1a**). TEV, on the other hand, does not exhibit any structural or conformational changes during the proteolysis of ENLYFQ/G and also does not possess IDRs(*33*) (**Fig. S1b**). We thus consider ADK as positive control and TEV as negative control.

We first compare the mechanical properties of the enzymes using diffusion as a proxy. These were determined using a microfluidic system consisting of two input channels that converge into a single output channel(*34*) (**Fig. 1b, see microfluidic setup section in methods**). We use the inlet 1 for the enzyme and the inlet 2 for the corresponding substrate (**see reaction scheme in Fig. S2**). Substrate concentrations were maintained at 10x of the respective Michaelis-Menten constant (*K_M_*). A pressure-driven system regulated the flow rates of both solutions in their respective channels. The enzymes were labeled with Fluorescein isothiocyanate (FITC) fluorescent dye and the fluorescence signal was measured near the outlet.

The experimental observations are shown in the lower panel of **Fig. 1b** and quantified in **Fig. 1c** to which we refer in what follows **(see estimation of fluorescence intensity shift section in methods).** The TEV stream shows no significant displacement towards the substrate flow (ii), while ADK makes significant inroads into the substrate stream (iii). On the other hand, comparing ADK (iii) and SrtA (iv), it is seen that SrtA exhibits greater diffusion into the substrate stream than ADK. We may correlate this with the conformational flexibility of the enzymes. In all these experiments, the conformational dynamics of the enzymes was not pre-activated. We now reduced the concentration gradient of the signal peptide between the streams by adding 160 µM -LPETGSSC- and 360 µM polyG in inlet 1, while maintaining substrate concentrations of 400 µM -LPETGSSC- and 360 µM polyG in inlet 2 as in the previous experiment (**Fig. 1b(iv)**). In this case, due to the addition of the substrates in inlet 1, the SrtA enzyme is already activated (**Fig. 1b(v)**) before the streams mix.

Since a gradient in the signal peptide -LPETGSSC-concentration should produce a diffusiophoretic flux, decreasing this gradient should lead to lesser diffusion. Additionally, if the mechanical dynamics of the enzymes did not affect their motion, the net downward flux of SrtA must not increase. However, we observe greater diffusion of SrtA (**Fig. 1c(v)**). This indicates that the pre-activation of SrtA is responsible for the increased diffusion, emphasizing that the mechanical activity of the enzyme plays a significant role.

The second important characteristic of SrtA is its ability to spontaneously phase separate into liquid droplets in solution at concentrations of 15 μM or higher (**Fig. S3a,** see **spontaneous phase separation of SrtA section in methods**). In contrast, neither ADK nor TEV exhibited phase separation, even at higher concentrations of up to 30 μM (**Fig. S3b, S3c**). We investigated the self-propulsion of SrtA within these liquid condensates while varying the reaction rate by adjusting the concentrations of the signal peptide [-LPETGSSC-] (**Fig. 1d**, **Fig. S4**). Importantly, the substrate is uniformly distributed within the droplets, which limits the potential for diffusiophoresis. Consequently, any variation in enzyme diffusion can be attributed to an enhanced conversion of chemical energy into mechanical activity. To monitor the propulsion of SrtA, we measured the diffusion rate of FITC-tagged enzymes using Fluorescence Recovery After Photobleaching (FRAP) with a confocal microscope. We observed more than a three-fold increase in the enzyme diffusion, from 0.21 ± 0.08 μm²/s at 0 μM to 0.73 ± 0.02 μm²/s at 400 μM of -LPETGSSC-, eventually reaching saturation (**Fig. 1d, see FRAP experiments section in methods**). The increase in motility of the enzymes is due to increased transpeptidation rate (upon increase of the substrate) and leads us to expect that the mechanical power of the SrtA can be effectively tuned by -LPETGSSC-concentration.

These findings underscore the unique properties of SrtA, particularly its ability to phase separate and self-propel. Somewhat surprisingly, even though there is increased motility of the enzymes, the liquid droplets of SrtA as a body did not exhibit any detectable motion at any substrate concentration.

### Synthesis of spherical vesicles and attachment of enzymes

We synthesized GUVs using a lipid mixture of DOPC and DGS-NTA(Ni) in a 97.5: 2.5 mass ratio (see **GUV preparation section in methods**). To stabilize the GUVs and their spherical shape, we maintained a pH7 buffer composition containing 350 mM sucrose, 3 mM CaCl_2_, 50mM HEPES, 50 mM KCl and 100 mM NaCl both inside and outside the GUVs. Enzymes are attached on the leaflets of the membrane using affinity-based interactions between the Ni^2+^ ions from DGS-NTA(Ni) lipids and 6x-His tags at the C-termini of the enzymes **(Fig. 2a)**.

**Fig. 2:**
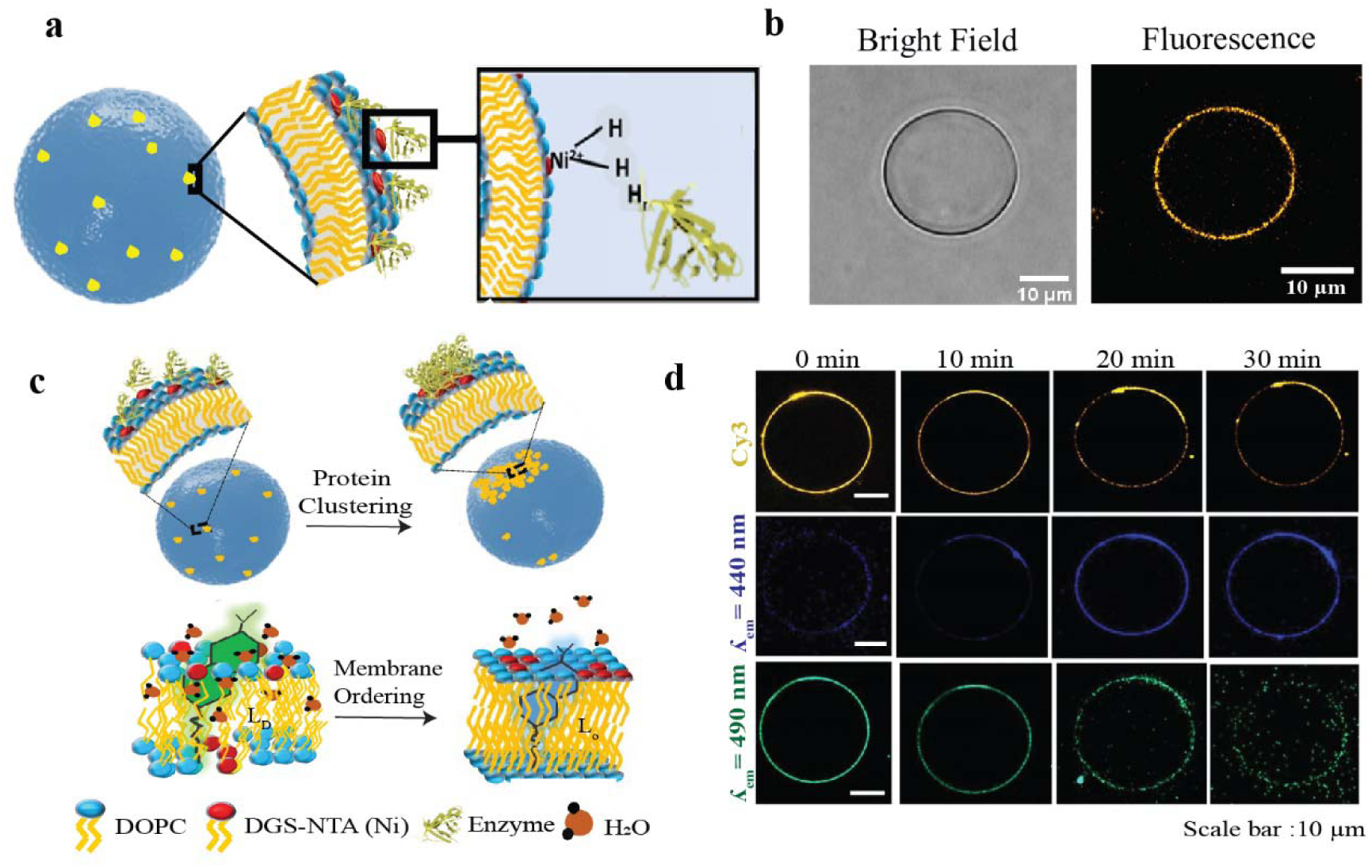
Spontaneous symmetry-breaking on membrane. **(a)** Schematic illustration of the enzyme attachment to the lipids of GUVs using 6xHis-Ni² affinity interactions. **(b)** Representative images of a spherical GUV captured in bright-field mode, alongside the enzyme-attached GUV displayed in fluorescence mode. The orange color in the fluorescence image indicates the presence of the Cy3-labeled enzyme. **(c)** Pictorial illustration depicting the clustering of SrtA enzymes on the GUV membrane (upper panel). Representative schemes show the emissions from the embedded Laurdan dye in the disordered (L_d_, left) and ordered (L_o_, right) states (lower panel) of the lipid membrane. **(d)** Fluorescence images of Cy3-tagged SrtA on the membrane, illustrating the clustering process of SrtA over time (upper panel). The middle panel displays the response of the Laurdan dye in the 440 nm emission channel, indicating lipid ordering (L_o_), while the lower panel shows the Laurdan dye response in the 490 nm emission channel, reflecting lipid disorder (L_d_). All images were captured using a confocal microscope.

Filtering the GUVs after passing through the Sephadex G-50 size-exclusion column allows us to isolate GUVs of a definite size. We study two groups of GUVs with diameters: 29 ± 6 µm as large GUVs (xGUVs) and 3.4 ± 1.6 µm as small GUVs (sGUVs), as determined from bright-field (BF) images (**Fig. S5**).

### Spontaneous Symmetry Breaking: Enzyme Clustering and Lipid Ordering on GUVs

While a primary reason to use GUVs as carriers is their biocompatibility, the fluidity of the lipids in the GUV membranes allows us to manipulate lipid organization and create clusters of reaction sites on the surface. This leads to breaking of the spherical symmetry of the GUV leading to directed propulsion. Using fluorescent tag (Cy3) on enzymes, we characterize their attachment and distribution on the GUVs (**Fig. 2b**). Initially, all three enzymes are seen to be uniformly distributed on the GUV membrane (**Fig. S6a**). However, the clustering propensity (**Fig. 2c)** of SrtA disrupts this uniform distribution (**Fig. S6a upper panel**) leading to the localization of fluorescence with time (**Fig. 2d upper panels**). Clustering is also demonstrated by using Laurdan dye which embeds into the membrane and emits light at 440 nm in ordered lipid phases (L_o_) while emitting at 490 nm in disordered phases (L_d_)(*35*) (**Fig. 2c lower panel, see membrane ordering induced by SrtA in Methods)**. The gradual clustering of SrtA on the membrane correlates with the progressive blue shift in the emission from the dye (**Fig. 2d, middle, lower panels**). No such clustering was observed for ADK and TEV (**Fig. S6a, middle and bottom panels**). This clustering of SrtA induces lipid ordering and denser packing with the expulsion of water in the membrane of the GUVs. A densely packed membrane will therefore limit any further mobility of SrtA on the GUV surface. Thus, upon photobleaching a SrtA cluster on the GUV surface, we observed a threefold decrease in fluorescence recovery compared to the diffusion of free lipids (**Fig. S6b**).

Previous studies have demonstrated that protein-induced lipid domain formation enhances membrane rigidity(*36*). Therefore, the arresting of diffusion taken together with the rigidity of the membrane will enhance the transfer of forces generated during folding/unfolding dynamics of the enzymes to the GUV, potentially resulting in observable translational and rotational motions of the entire structure.

### Dynamics of GUVs

Given the clustering and immobility of the enzymes, we anticipate a certain degree of cooperativity during the folding and unfolding reaction cycle, attributed to steric and proximity effects. Consequently, we considered the resultant forces (***fi***) generated by the various enzyme clusters and represented the model GUV schematically in Fig. S7. Since these clusters do not diffuse, the forces remain anchored to fixed points on the GUV surface. Therefore, we can consolidate these contributions into a single effective radial force ***F*** acting at a fixed surface point P. The tangential components of these forces will result in an effective torque τ. As the GUV rotates, the point of application P of the effective force also rotates. Generally, the effective force ***F*** will not lie in the plane of the torque τ, leading to helical trajectories, provided that the gravitational component responsible for the sedimentation of the particle is dominated by self-propelling forces. Such trajectories of particles in the Stokes regime are described by the chiral Active Brownian Particle (cABP) model (**See Tracking of sGUVs in Methods**)(*8*, *37–39*). Later, we will use cABP model to quantify the dynamics of sGUVs.

We initiate transpeptidation by introducing the reactants, -LPETGSSC- and polyG, into the solution containing SrtA-coated GUVs. BF and wide-field fluorescence microscopy were employed to monitor the GUVs and their activity. We designed a closed chamber for our experiments (**Fig. S8**) and ensured the absence of hydrodynamic artifacts in our measurements as described in Methods (**See Sample Chamber for Imaging in Methods).** To monitor the activity of GUVs, we used the Nile red fluorophore that is adsolubilized in lipids and illuminates the membrane as red. After initiation of transpeptidation, post a transient period, we observed dynamic fluorescent patches on xGUVs that appeared to rotate (**Fig. S9, Movie S1**).

To quantify the rotational trajectories of xGUVs, we introduced small lipid vesicles (tracers) composed entirely of DOPC (diameter less than 3 μm) into the solution. These tracers bind non-specifically to the xGUV membrane, allowing us to track their trajectories. The number of tracers was controlled for 1:1 binding. From the BF imaging, we found that nearly 67% xGUV were bound to single tracers, while around 4% were attached to two tracers. Of the remaining xGUVs, nearly 14% did not have any tracers attached, and about 15% ruptured (**Fig. S10**). Because of their small sizes, the trajectories of sGUVs were quantified using the cABP model **(See Tracking of sGUVs in methods).**

Upon activation, the GUVs exhibited a variety of motion which depends on their sizes as we will now describe.

### Active spinning of xGUVs (d > 20 **µ**m)

In sharp contrast to existing literature, nearly all the xGUVs exhibited uninterrupted, unidirectional spinning (**Movie S2**). The spinning continues for **hours** (**Fig. 3a**) until the xGUVs actually ruptured. The spin axis of all xGUVs were observed to be perpendicular to the slide surface. However, the center of mass (CoM) of the xGUVs exhibit negligible translation (**Fig. 3a upper panel**). A small number with relatively smaller diameters (d < 25 µm) displayed transient dynamics, where the gyrations stabilized after about 3 minutes (**Movie S3, Fig. 3a lower panel**). The jitter in the trajectory shown in this figure is due to fluctuations in the position of the tracer particle.

**Fig. 3:**
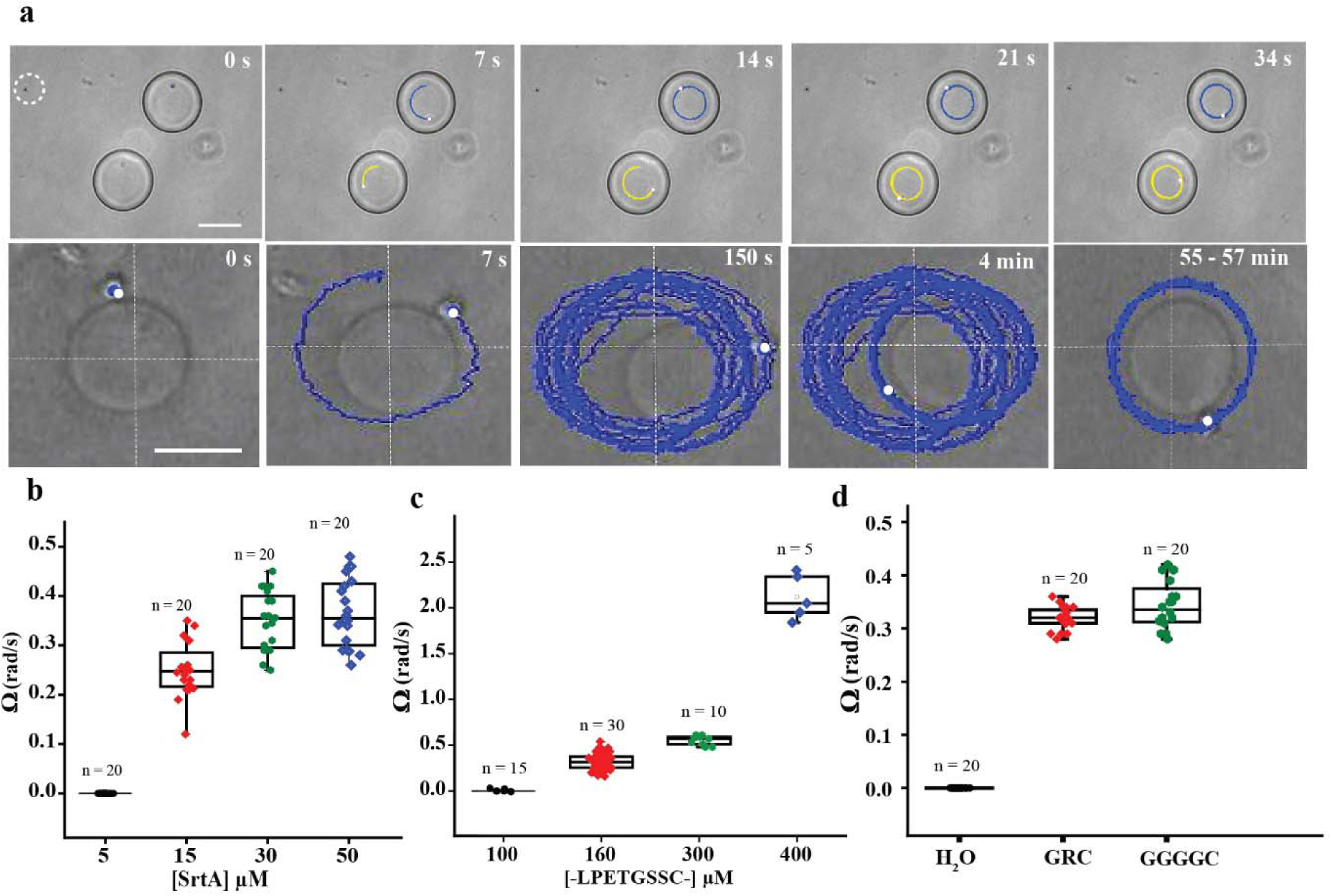
Active rotation of SrtA embedded xGUVs. **(a)** The upper panel displays snapshots of two unidirectional spinning xGUV with a single tracer each at the core. An immobile tracer particle (white circle) in the left upper corner can be taken as control. The lower panel displays the snapshots of a xGUV of diameter (∼25 µm) with a tracer on the periphery, exhibiting gyrations for up to 150 seconds before settling to pure spinning. Reference axes are shown as dotted white lines to track the CoM. Scale bars: 20 µm **(b)** Scatter plot showing the variation in angular velocities with concentration of SrtA. **(c)** Scatter plot illustrating the change in angular velocities of xGUVs with the concentration of -LPETGSSC-. **(d)** Scatter plot depicting the variations in angular velocities with different nucleophiles. ‘n’ mentioned in the figures (b - e) refers to the total number of GUVs collected from 3-4 different experiments.

We performed the experiments four times and obtained approximately 73% spinning xGUVs. Of the remaining, around 14% of xGUVs showed no visible tracer attachment, while nearly 13% were either stationary or damaged. By tracking the position of the tracers, we determined the angular velocity (Ω) of the xGUVs.

The angular velocity of the xGUVs can be controlled by the enzyme loading. A change of SrtA concentration from 15 µM to 50 µM on xGUVs leads to an increase in Ω from *0.25 ± 0.05 rad/s* to *0.36 ± 0.06 rad/s* (**Fig. 3b**). Beyond 50 µM, xGUVs distort significantly and aggregate (**Fig. S11a**). Similarly, increasing the fraction of SrtA by incorporating more than 2.5% DGS-NTA(Ni) also induced distortions in the xGUVs (**Fig. S11b**). At a concentration of 5 µM SrtA, where the enzyme did not form droplets in solution, xGUVs remained inactive. All experiments described henceforth use 50 µM of SrtA unless specified otherwise.

SrtA executes transpeptidation in two consecutive steps, a rate-determining thioesterification on signal peptide (-LPETGSSC-) with conformational and structural changes, followed by a nucleophilic attack from the primary amine of Gly (**Fig. S2a**) which reverses the physical changes in SrtA. The large differences in the reaction rates implies that this cyclic process is non-reciprocal (*40*). We can thus change the reaction rates by independently altering the concentration of -LPETGSSC- or the chemical nature of the nucleophiles in solution (**Fig. S12, Fig. S13**). We first increased the concentration of -LPETGSSC- in solution, and observed an increase in Ω from *0.32 ± 0.08 rad/s* at 160 µM to *2.1 ± 0.23* rad/s at 400 µM (**Fig. 3c, Movie S4**). This is in line with the increasing self-propulsion of SrtA in liquid droplets with - LPETGSSC- (**Fig. 1d, S4**). Beyond 400 µM concentration of -LPETGSSC-, the angular velocity becomes large enough to eject the tracer particles (**Movie S4**). Overall, these results suggest one method to generate more mechanical energy and hence more speed.

We next altered the rate of the second step in the reaction scheme (**Fig. S2a**) by varying the nucleophiles, H_2_O, -GRC-, and polyG, with increasing affinity towards the thioester. Among these three, the rate of hydrolysis of the thioester is the slowest, and the rates of nucleophilic attack for -GRC- and polyG are comparable (**Fig. S13**). We observed a significant impact of nucleophiles on xGUV spinning. With H_2_O as the nucleophile, xGUVs remained stationary (**Fig. 3d**). We hypothesize that this is due to the slowing in the rate of the second step (**Fig. S2a**) which essentially reduces the non-reciprocity of the cyclic folding/unfolding of the enzyme. Comparing -GRC- and polyG as nucleophiles, the spin frequencies of xGUVs are similar (**Fig. 3d**).

A striking feature of the xGUV motion is the persistence of the rotation. The GUVs continued to spin without pausing until they ruptured. We observed minimal variation in the angular speed (Ω) over a 60-minute observation period (**Fig. S14**). Results in **Fig. S15** shows that the product (- LPETGGGGC-) alone can perform the entire catalytic cycle of SrtA without the need for additional reactants (-LPETGSSC- and polyG). We attribute the sustained spinning to this apparent cyclic nature of the reaction. However, the product and/or the reactants are eventually consumed via hydrolysis which is many-fold slower than the transpeptidation (**Fig.S2a**).

The dynamics of the xGUVs coated with SrtA contrasts sharply with the controls that contain either ADK or TEV which entirely failed to spin or translate during catalysis (**Fig. S16**). Based on the microfluidics experiments, the TEV-coated xGUVs were expected to remain inert, but not the ADK-coated xGUVs. We hypothesize that the lack of clustering and the absence of lipid ordering, which are typically associated with increased membrane rigidity, are responsible for the inactivity of ADK-coated xGUVs. We were, however, unable to induce clustering of ADK on large GUVs. As we demonstrate in a subsequent section, forcing ADK to cluster and lipids to order on sGUVs does indeed induce active motion.

We note here that, in rare cases, the spinning of the xGUVs is intermittent (**Movie S5a)**. From **Fig.S7**, we can see that the activation/deactivation of any cluster will lead to a change in the force and torque. Alternatively, the change in direction could have arisen from the application of large random force/torque.

Before activation, the xGUVs settled onto the microscope slide, as confirmed by the observation that their sedimentation plane aligns with that of a heavy magnetic bead evinced by the fact that the two come into focus together under observation (**Movie S6**). It is important to note that, for these xGUVs, the self-propulsion forces were insufficient to produce any translational movement **(See Sedimentation Estimates for GUVs in Methods**). It is plausible that due to its own weight, the xGUVs undergo deformations. In fact, in our measurements of lipid diffusion at the poles (bottom and top) of the xGUVs using FRAP, we observed relatively slower diffusion at the bottom pole (effective diffusion coefficient, *D_eff_* = 0.08±0.01 μm^2^/s) compared to the top pole (*D_eff_* = 0.30 ± 0.09 μm^2^/s) **(Fig. S17; See FRAP experiments in Methods)**. While the xGUVs settle on the surface, they nevertheless can move or roll under large external perturbations (**Movie S6**).

### Gyration of sGUVs

Unlike the xGUVs, SrtA embedded sGUVs display a variety of rotation, translation, and gyrations (**Movie S7, Fig. 4a**). We emphasize that the length and time scales of motion depicted in these figures are unprecedented. The smoothness of the trajectories also makes it clear that the motion is not diffusive in the observation time scale. Our samples are dominated by those sGUVs which remain within the 300 μm field of view during the imaging (many sGUVs move over much longer distances as seen in **Movie S7**).

**Fig. 4:**
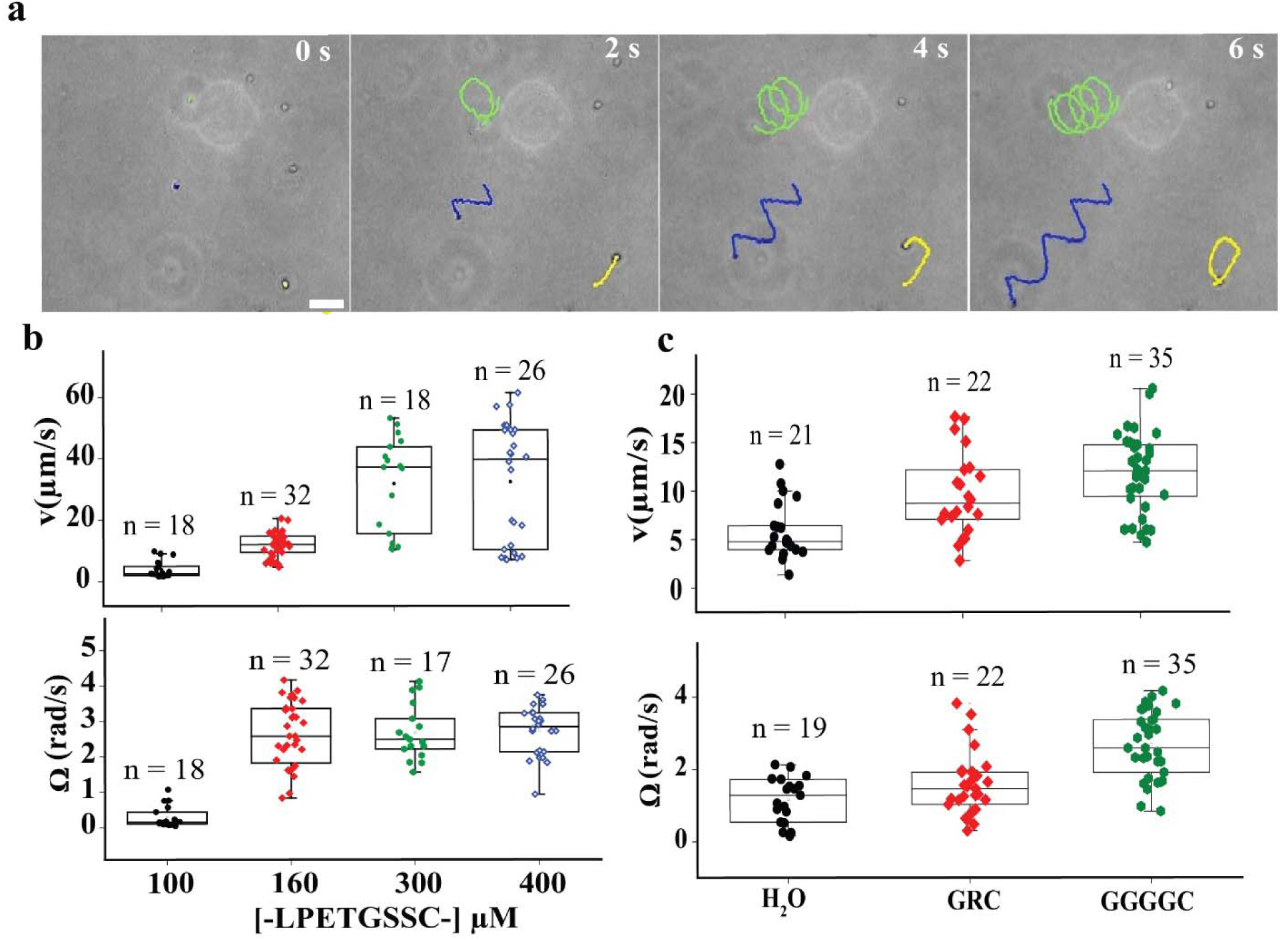
Varied Motion of SrtA embedded sGUVs. **(a)** Snapshots illustrate the simultaneous rotation and translation of SrtA embedded sGUVs. The yellow curve tracks the motion of an in-plane sGUV, while the green and blue lines track out-of-plane helical trajectories. Scale bar: 10 µm. **(b)** The trend in linear speed (v) and angular velocity (Ω) with -LPETGSSC-concentration. **(c)** Variation in v and Ω of sGUVs with different nucleophiles. ‘n’ in the figures (b - c) refers to the number of sGUVs collected from 3-4 different experiments.

In many cases, the motion of the sGUVs was planar with the planes randomly oriented. We also observed sGUVs that gyrate in three dimensions with helical trajectories as expected from Newton’s laws of motion. As with the intermittent motion of xGUVs, we observed occasional shifts in the direction of motion for sGUVs (**Movie S5b, Movie S7**). While inactive sGUVs eventually settle on the slide, the downward drift due to gravity is more than counteracted by the self-propelling forces (**See Sedimentation Estimates for GUVs**).

We quantified the motion of those sGUVs that remained in the focal plane of the imaging set-up. While this introduces a sample bias, it allows us to analyze the trajectories accurately and does not affect any of the conclusions below. From these trajectories **(Fig.4a)**, we determined the velocity autocorrelations (VACF) and the mean square displacements (MSD) **(See Tracking of sGUVs in Methods, Fig. S18).** VACF determined from the trajectories, showed decaying oscillations (**Fig S18a**). The MSDs showed a superdiffusive regime with MSD (t) ∼ t^V^ (v> 1), an intermediate oscillatory regime (a crucial signature of a cABP), and subsequent diffusive regime with MSD (t) ∼ t (**Fig S18b**). Fitting these experimental VACF and MSDs with the corresponding analytical expressions of a cABP **(See Trajectory Analysis in** Methods) shows excellent agreement and allows us to determine the effective self-propulsion speed of the SrtA coated sGUVs (v) to be 11.9 ± 4.0 µm/s and angular speed (Ω) of 3.0 ± 1.3 rad/s. The goodness of the fit of the cABP model also provides additional indirect evidence that clustering leads to point forces that propel the sGUVs (**Fig.S18**). It may be mentioned that a direct estimation of the angular and linear speeds is possible for a few GUVs that carry a distinguishing mark on their surface. The values so obtained are comparable to those obtained from the above fit.

We will next discuss the effects of varying the concentration of -LPETGSSC-, which leads to a variation in the reaction rates **(Movie S8)**. Unlike the xGUVs, even at 100 µM of -LPETGSSC-, the sGUVs showed dynamical motion. The self-propulsion speed, *v* of sGUVs appears to saturate at 300 µM of -LPETGSSC- **(Fig. 4b)**. More strikingly, Ω saturated at 160 µM concentration of the peptide (**Fig. 4b**) where the xGUVs were just beginning to spin. It is reasonable to anticipate that GUVs with higher curvature exhibit a stronger affinity for the substrate (*41*), leading to saturation at lower concentrations of -LPETGSSC-.

The observations with nucleophiles were also different for sGUVs compared to the xGUVs **(Fig. 4c)**. sGUVs showed both linear (*v* = 5.51 ± 2.47 µm/s) and rotational (Ω = 1.17 ± 0.63 rad /s) motion even with hydrolysis, though the magnitudes are significantly slower than with polyG. Between -GRC- and polyG, we observed comparable v and Ω values, an observation similar to xGUVs. Further, with H_2_O as a nucleophile, we observed a gradual drop in the activity of sGUVs with time to Brownian motion.

### Design Principles

Based on our experiments detailed above, we may assemble the following ingredients that seem to be implicated in generating actively propelled, spinning GUVs.

The first element required is a flexible active enzyme which undergoes a cyclic but non-reciprocal conformation change. The latter feature ensures the violation of time reversal symmetry as required by the Scallop theorem (*40*). In our case, TEV which may be termed inflexible does not generate active dynamics in any circumstances, whether in solution (Microfluidics) or on the GUV surface. In the hydrolysis route, the difference in the rates of the nucleophilic attack and the transpeptidation is decreased. This effectively decreases the asymmetry in the folding and unfolding rates which may be the underlying reason for the reduced activity.

The second design element is clustering of the enzymes. We note that GUVs coated with either ADK or TEV do not exhibit any active dynamics **(Fig. S19)**. Since ADK and TEV do not phase-separate, the distribution of ADK and TEV on the surface of the GUVs is uniform **(Fig. S6a)** leading to pure diffusion of the GUVs. Therefore, we induce clustering by using a third saturated lipid, DSPC, on the GUVs **(See GUVs preparation section in methods**). DSPC is known for its propensity to undergo phase separation, forcing the lipids to pack densely and consequent rigidity of membrane(*42*) (**Fig. S20)**. In ADK embedded sGUVs with a 1:1 distribution of DOPC and DSPC, approximately one hemisphere is covered with enzymes. We recovered active dynamics with a v of *3.07 ± 2.25* µm/s and Ω of *0.21 ± 0.16* rad/s. Increasing the DSPC fraction increased the magnitude of v and Ω non-linearly (**Fig. 5a-f**) and was maximum at 100% with v *of 13.32 ± 4.22* µm/s and Ω *of 3.14 ± 1.75* rad/s, respectively (**Movie S9**). These values are comparable to those of SrtA-embedded sGUVs at any DSPC concentration. Correspondingly, the velocity autocorrelation and MSD data for ADK with 100% DSPC compared well with that of SrtA-embedded sGUVs **(Fig. S21, S22)**. No motility was observed in sGUVs with TEV, irrespective of DSPC content. SrtA coated GUVs showed no significant changes in activity with increasing DSPC content (**Fig. 5g-5l**). We attribute this to the spontaneous phase-separation behavior of SrtA, which caused the enzymes to segregate from the get go. Additionally, with 100% DSPC, we observed ta large spread in both v and Ω. In this case though, we note that increasing DPSC concentration appears to destabilize the sGUVs.

**Fig. 5:**
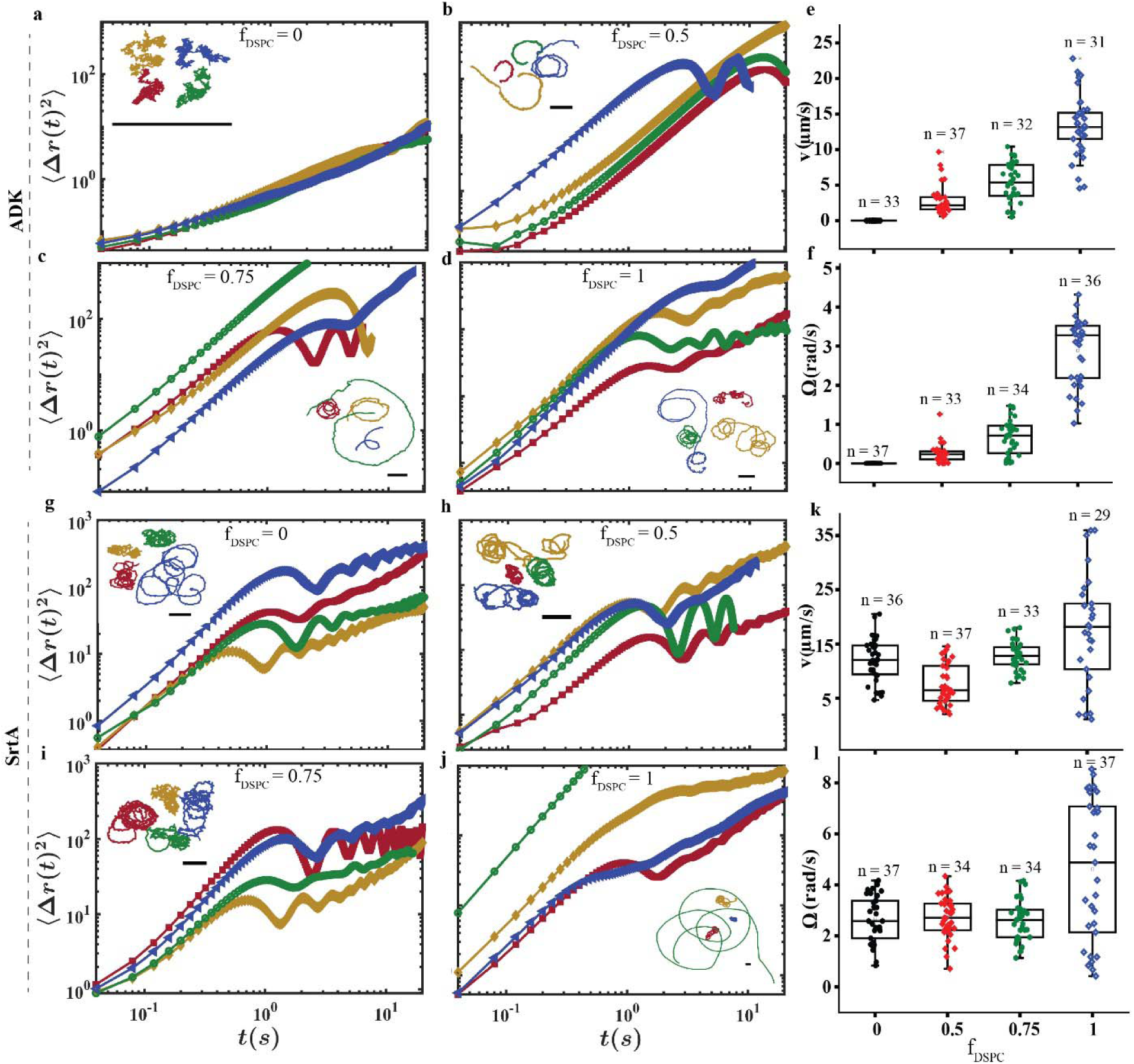
Enzyme Clustering and Active Dynamics in sGUVs. **(a-d)** Representative MSDs of ADK coated sGUVs are shown at increasing fractions of DSPC 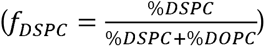:(a) 0, (b) 0.5, (c) 0.75, and (d) 1. The insets display the corresponding trajectories of the sGUVs. Scale bar: 10 μm. **(e) & (f)** show the trend in v and angular velocity, respectively, with increasing f_DSPC_ for ADK-coated sGUVs. **(g-j)** MSD plots for SrtA-coated sGUVs with increasing f_DSPC_ from left to right: (g) 0, (h) 0.5, (i) 0.75, and (j) 1. **(k) & (l)** show the corresponding trend in v and Ω, respectively. ‘n’ in the figures (i, f, k, and l) refers to the number of GUVs collected from 3-4 different experiments.

The final design element required can be termed membrane rigidity. Activation of the enzymes leads to larger forces (diffuse more rapidly), but arresting diffusion can transfer the force to the GUV, only if the chassis material is rigid. In our case, this rigidity is achieved by forcing lipid ordering. This is supported by the observation that SrtA in liquid droplets is mechanically active as confirmed by FRAP **(Fig.1d, S4**), but does not lead to motion of the droplets. On other hand, GUVs coated with ADK do not induce lipid ordering and hence the GUVs were not rigid, so were not able to rotate or translate. GUVs coated with SrtA were always dynamic for reasons already discussed.

We can add two further elements. The *speeds* of the GUVs can be manipulated by altering the rates of the folding/unfolding reaction of the enzymes. As we have shown, the angular and linear speeds can be increased several folds by increasing the peptide concentration **(Fig.3c, 4b)**. Modifying the nucleophiles can also produce a similar effect **(Fig.3d, 4c)**.

Finally, we must mention that the very long *duration* of the GUV dynamics in our case depends on the apparent cyclicity of the reaction scheme **(Fig.S15)**. In other cases, this may be manipulated by the variation in concentrations and the suppression of dissipation routes such as hydrolysis.

## Summary and Discussion

In summary, we proposed that *clusters* of *flexible* enzymes distributed anisotropically on any *rigid* object can be used to generate a variety of motion, from pure rotation to complex gyrations. The magnitudes of angular and linear speeds would depend on the number of clusters and the extent of asymmetry in the distribution of the clusters. In fact, we have also conducted experiments to verify that preventing the SrtA from clustering on the GUVs completely disrupts their motion. This is, of course, consistent with the observation that inducing clustering of ADK enzymes leads to chiral motion **(Fig. 5a-5f)**. In our case, the apparent cyclicity of the enzyme reaction produced very long durations of dynamics. Surprisingly, the magnitudes of linear speeds and rotational frequencies were not highly sensitive to the enzymes’ powering abilities with ADK and SrtA producing similar speeds **(Fig. 5e-f, 5k-l)**.

Thus, our work should pave the way for the development of a variety of artificial systems capable of mimicking natural gyrotactic behavior and opens new avenues for targeted drug delivery, minimally invasive surgery, and diagnostic tools. Being free from the influence of directional biases, our spherical µ-rotors offer a unique platform for studying emergent behaviors.

There are several important questions which our experiments were unable to completely resolve. We have presented evidence that can be interpreted to mean that the motive power arises from the mechanical action of the enzymes rather than diffusiophoresis **(Fig.1b, c)**. It is clearly important to understand the mechanism that is at the root of the dynamics we have shown. An improved set of microfluidic experiments can be performed to differentiate the diffusiophoretic contributions from mechanical.

An important feature that remains puzzling is the complete absence of translation in the xGUVs observed. From the force estimates **(See sedimentation section in material method)**, one could expect translation or at least rolling motion. It is plausible that the xGUVs deform slightly when they settle on the slide preventing rolling **(Fig.S17)**. However, coating the surface of the slides with PEG did not produce any significant translation in the xGUVs either.

In our experiments, clustering, and rigidity (lipid ordering) occur together because of SrtA. It will be interesting to quantify and manipulate the amount of rigidity induced by the clustering and it would be desirable to study these effects separately. And conversely, one can coat similarly clustering enzymes on rigid objects such as nanoparticles, polystyrene beads, silica beads. One can engineer more buoyant spheres by filling the inside with some lighter substance and may achieve gyration even with large particles.

Using the Stokes equations, we estimate the F of ∼ 0.4 – 3 pN for GUVs of varying sizes. On the other hand, individual motor proteins typically generate forces of a few pN. In sharp contrast, the entire cluster of around 10^4^-10^6^ enzymes in our experiments seem to produce only a few tenths of a pN. Future studies could test these hypotheses by inducing *ordered* clustering of enzymes. This could lead to cooperative force generation and remarkably large velocities of the entire GUV assembly.

However, it has been pointed out that for Volvox, there is significant dissipation that should be expected because the point of application of the forces is a distance ER (0<E<<1) away from the surface (*43*). In fact, translation forces pick up a factor E^2^ while the torques are suppressed by single factor of E. Given that, E ∼ 10^-3^, this could be the reason for the small magnitudes compared to motor proteins and also perhaps explain the absence of translation in the large GUVs.

We conclude by making one further surprising observation. Our observations show that v and Ω decrease with increasing particle size, similar to that observed in Volvox(*3*, *4*) and synthetic active particles (*44*). Such behaviour has been attributed to increased hydrodynamic interactions and resistance in the fluid. The extrapolated values of the shear stress obtained for Volvox to smaller radii are comparable to the shear stresses generated by our artificial μ-rotors **(See Fig. S23 and the figure legends for estimate)**. These observations could suggest an underlying scaling principle which merits further exploration.

The literature on microorganisms includes several studies showing a variety of cooperative behaviour when these objects are in proximity. In our experiments as well, we have observed a few instances of “attractive” interaction between two sGUVs leading to intermittent pairing **(Movie S10)**. Another striking observation which may well be a consequence of such cooperative interactions is that nearly all the xGUVs in a given observation were seen to spin in the same direction. Across observations though, this direction occurred with either handedness. A future objective would thus be to introduce a control over the handedness of rotation.

## Acknowledgments

We sincerely thank Mr. S.M. Rose and Prof. Dr. Sharmistha Sinha (Institute of Nanoscience and Technology, India) for helping us with plate-reader; Ms. Priyanka and Prof. Subhabrata Maiti (IISER Mohali) for helping us with the Microfluidics Experiments; Mr. Souvik Halder (NISER Bhubaneswar) for helping us in deciphering the cyclicity of SrtA reaction as a part of summer dissertation under the guidance of VK & SR; Prof. Anil K Dasanna for critical reading and comments; Ms. Anweshika Pattanayak for sharing knowledge on the cABP model.

We also thank ‘perplexity.ai’ and ‘ChatGPT’ for assisting in editing draft in some cases for a better reading.

## List of Supplementary Materials

Materials and Methods Supplementary Text

Figs. S1 to S23 References (*26–32*) Movie S1 to S10

## Materials and Methods

### Expression and purification of Enzymes

SrtA(Δ59) (Addgene plasmid #51138), TEV Protease, and Adenylate kinase (Addgene plasmid #73625), all tagged with 6x His, were expressed in E. coli BL21 (DE3) or Rosetta cells. For SrtA, cells were grown at 37°C until OD600 reached 0.5, induced with 0.5 mM IPTG, and incubated at 25°C for 16 hours. TEV Protease was expressed by growing cells at 37°C until OD600 reached 0.4, induced with 0.3 mM IPTG, and incubated at 16°C for 20 hours. Adenylate kinase was grown at 37°C until OD600 reached 0.6, induced with 1 mM IPTG, and incubated at 37°C for 4 hours. Cell pellets were resuspended in a specific lysis buffer, lysed by sonication, and the lysates were centrifuged. The supernatants were passed through pre-equilibrated Ni-NTA columns, and the enzymes were eluted using 250-300 mM imidazole. The purity of the enzymes was verified by SDS-PAGE, and the enzymes were stored at -80°C.

### Fluorescence Tagging of Enzymes

A stock solution of FITC (1 mg/mL) was prepared in DMSO. Next, 100 µl of the FITC stock was added to 60 µl of 60 µM of enzymes to maintain a [dye]:[enzyme] ratio of 1:10. The solution was incubated in the dark at 4°C for 7 hours to allow for the tagging reaction to occur. Subsequently, 50 mM ammonium chloride was added to quench the ongoing reaction, and the mixture was allowed to incubate for an additional 2 hours. The reaction mixture was then passed through a Sephadex G-25 column, and 200 µl fractions were collected. UV absorption spectra were recorded for each elution, revealing that the conjugate of FITC and the enzyme displayed an absorption maximum at 280 nm, while FITC exhibited a peak at 495 nm. We compared the absorbance maxima and observed an enzyme to dye ratio of 1:3.

### Microfluidic setup and Estimation of fluorescence intensity shift

A microfluidic channel with two inlets and one outlet, measuring 17 mm × 0.6 mm × 0.1 mm (L × W × H), was utilized. Enzyme labeled with FITC was injected from one inlet, while the corresponding enzyme substrates were introduced from the other inlet at a flow rate of 0.3 ml/h.

Images were captured 16 mm downstream from the point where the inlets merged, using a fluorescence microscope (Zeiss Axis Observer 7 microscope).

Fluorescence intensity profiles were plotted at the outlet across the channel width (600 µm). The profiles were normalized to the maximum intensity observed. Notably, an apparent drift of SrtA and ADK towards the substrate channel was recorded. Control experiments were conducted without substrate in the buffer in the second channel, revealing a normalized intensity of 0.5 near the channel midpoint, specifically at 300 µm. In contrast, when comparing the same for SrtA/ADK in one channel and substrates in the second channel, we found that the normalized fluorescence of 0.5 had spatially shifted from 300 µm to higher values. A greater spatial shift correlates with a higher drift, indicating increased propulsion.

Throughout each measurement, we ensured that the two flows at the inlets converged almost precisely at the channel’s centerline. Importantly, the interface between the flows at the inlets deviated by less than 3 µm from the centerline. This deviation is significantly smaller than the observed drift near the outlet (1.6 mm downstream) in the SrtA/ADK with substrate experiments, as confirmed across multiple experimental trials.

### Spontaneous Liquid-Liquid Phase Separation (LLPS) of SrtA

Purified recombinant SrtA was stocked at approximately 400 µM in a buffer containing 50 mM HEPES, 100 mM NaCl, 50 mM KCl, and 3 mM CaCl2 (pH 7.5). Before conducting the phase separation assay, the sample was centrifuged at 15,000 rpm for 20 minutes at 4°C to eliminate any aggregates. To induce phase separation, the concentrated sortase solution was diluted to various concentrations. Notably, at concentrations of 15 µM and above, we began to observe droplet formation. The formation of liquid droplets was monitored using a brightfield microscope and a fluorescence confocal microscope (ZEISS LSM 980 Elyra 7) for approximately 20 minutes, during which new droplets formed and fused with existing ones. After 20 minutes, we measured a distribution of droplet diameters of approximately 1 µm ± 0.8 µm.

A similar procedure was employed for the adenylate kinase (ADK) and TEV protease proteins; however, no droplet formation was observed even at enzyme concentrations of up to 50 µM.

### FRAP experiments

FRAP (Fluorescence Recovery After Photobleaching) experiments were conducted using a ZEISS LSM 980 Elyra 7 confocal microscope. The FRAP experiments on sortase A (SrtA) droplets were performed under two conditions: with and without substrate. We systematically varied the concentration of the substrate -LPETGSSC- and conducted FRAP analyses. For all conditions, FRAP was performed on mature droplets with a diameter of at least 1.8 µm.

An optical beam area of 1 µm² within the SrtA droplets was photobleached using a 488 nm laser at a power of 500 mW for 5 seconds. The recovery of fluorescence after bleaching was recorded using Zen Blue 3.2 software (ZEISS). To assess the heterogeneity in enzyme activity within the droplets, we repeated FRAP measurements on three different droplets for each condition. The fluorescence recovery of the photobleached area within the droplet was monitored over time and normalized to the pre-bleach intensity. The fluorescence intensity profiles for the entire trace (pre- and post-bleach) were adjusted by subtracting the background fluorescent intensity. Data analysis and plotting were performed using Origin software.

For FRAP experiments on membranes, enzymes anchored to GUVs were subjected to photobleaching while maintaining all other experimental conditions consistent with those used for droplets. For the FRAP measurements at the poles of xGUVs, FITC-coated TEV proteins were utilized.

### FRAP data analysis

We estimated the recovery lifetime from the fitting of the fluorescence intensity recovery with time to following equation:

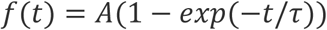

In above equation, A refers to mobile fraction of proteins and *τ* is the lifetime of the recovery. From the *τ*, we next estimated the half-life (*τ*_1/2_) of the recovery followed by the two-dimensional diffusion coefficient (D_eff_) as following:

*τ*_1/2_ = ln(0.5) /-*τ* and D_eff_ = (0.88*w^2^)/4*τ*_1/2_, where w is the radius of the region selected to study FRAP (bleach area).

### GUV Preparation

We utilized two sets of giant unilamellar vesicles (GUVs) with differing lipid compositions. The lipid compositions for xGUVs and a subset of sGUVs comprised 2.5% DGS-NTA (Ni) (790404 C, Sigma-Aldrich) and 97.5% DOPC (850375 P) or DSPC (850365 P, Sigma-Aldrich). To illustrate the design principle for spherical *μ*-rotors, we systematically substituted DOPC with a third lipid, DSPC (850365 C, Sigma-Aldrich).

GUVs were prepared using a poly(vinyl alcohol) (PVA) gel-assisted swelling method. Initially, glass slides were cleaned with piranha solution (a 3:1 mixture of H_2_SO_4_ and 30% H_2_O_2_) and subsequently coated with PVA, which was dried at 65°C. Following this, the slides were coated with the lipid mixture and placed in a vacuum oven until the solvent had completely evaporated. GUVs were then collected in a solution containing 350 mM sucrose and 50 µM of respective enzymes. The rationale to add enzymes during GUV harvesting is to maintain an isosmotic condition and thus, avoid any shape-deformation in GUVs. However, this results in the GUVs filling with protein, making them heavier. After harvesting, we pass the GUVs through Sephadex G-50 (Sigma) size-exclusion column and isolate them based on size. The formation of GUVs was visualized using phase contrast and fluorescence microscopy (Leica, DMi8). After isolation, GUVs are incubated with enzymes again to replenish any loss of outer surface proteins during the isolation process. Excess enzyme was removed with centrifugation at 13400 rpm for 60 s.

### Membrane Ordering induced by SrtA

Segregation of SrtA on the GUV membrane induces lipid ordering. We examined the local environment of the membrane lipids using Laurdan dye, which embeds into the membrane and responds to dipolar relaxations of water. In the case of ordered lipid phases (L_o_), Laurdan emits at 440 nm, whereas it emits at 490 nm for the disordered phase (L_d_). This shift is because Laurdan’s emission spectrum is sensitive to the polarity of its environment. In ordered phases (L_o_), the lipid molecules are tightly packed, expelling the water molecules. This results in a more non-polar environment for Laurdan, leading to a blue shift in its emission. Conversely, in disordered phases (L_d_), the lipid molecules are less ordered, allowing for more water molecule movement. This creates a more polar environment for Laurdan, resulting in a red shift in its emission.

We thus incubated GUVs with Laurdan dye (10 µM) for 30 min at room temperature for incorporation of dyes into the membrane and monitored the dye emissions by exciting the GUV containing dyes at 350 nm.

### Sample Chamber for Imaging

All experiments related to tracking active particles were performed under Brightfield-field inverted microscope and using a custom-made closed chamber. For the base of the chamber, we used microscope slides measuring 60 mm × 24 mm × 0.13 mm for LxBxH. Reusable press-to-seal silicone chambers, with a width of 0.8 mm and a diameter of 15 mm, were utilized as wells for the samples. A 22 mm diameter coverslip was placed on top to seal the chamber **(Fig S8)**. A 5*μ*L extract is pipetted out onto the microscope slides and the substrates added and the chamber is sealed with the coverslip. After a wait time of approximately half an hour, the enzyme-coated GUVs become activated and display a range of motions. In the case of xGUVs, we occasionally observed a brief jerk before they began to rotate in a steady manner.

Following this activation period, the GUVs exhibit steady motion which are recorded using a microscope with a field of view of about 300 **μ**m. The absence of hydrodynamic artifacts was verified by several control experiments described below. Passive polystyrene beads (2 µm) were suspended in a solution containing **inactive** GUVs. The absence of movement of the polystyrene beads corresponded with the GUVs remaining static. We also verified external flows such as a leak led to identifiable disruptions in the trajectories of the GUVs. During experiments involving TEV and unclustered ADK, no flow or active motion was detected in the GUVs, despite catalytic turnover, indicating the absence of external flows. Additionally, to mitigate any heating effects from the white lamp (light source) used in the microscope, we varied the power of the lamp and observed no visible impact on GUV activity.

### Reaction Kinetics of SrtA in Solution with varying substrates

SrtA undergoes catalytic turnover in two sequential steps, a thioesterification step with peptide containing -LPETGSSC-sequence followed by nucleophilic attack, specifically from the primary amine of Gly. To monitor the effect of the substrates in the kinetics, we first varied the concentration of the signal peptide (-LPETGSSC-) substrate systematically while keeping the nucleophile -GGGGC-fixed at 360 µM. The enzyme concentration was fixed to 15 µM. To monitor the progress of the reaction, we chemically modified the -LPETGSSC-substrate with a DNP quencher and a MCA fluorophore at the C-terminal and N-terminal, respectively. The final substrate construct becomes-MCA-LPETGSSC-DNP which is non-fluorescent. Cleavage of the T-G bond in the substrate would generate fluorescence at 400 nm when excited at 322 nm. We then monitored the kinetics from the increasing fluorescent intensity using a Plate-reader (Tecan Infinite Mplex) for 90 mins till we reached saturation. We performed the experiments in a 96-well plate with wells containing varying concentrations, 100µM, 160 µM, 300µM, and 400µM of MCA-LPETGSSC-DNP. Each experimental condition was repeated at least three times. The data presented in **Fig.S12** are the average data with standard errors for the precision.

Next, we modulated the second step by varying the nucleophiles (-GRC-, polyG, and H_2_O) and monitored the progress in reaction as described in the previous section. For each nucleophile, we performed the experiments at 4 different concentrations and estimated the maximum reaction velocity (*V*_max_) and Michaelis constant (*K_M_*) of Michaelis-Menten equation by following the already reported protocol (**Fig.S13**) (*40*).

### Sedimentation Estimates for GUVs

#### (a) sGUVs (∼4 µm diameter)

*Area of the sGUVs: A* = 4*π*(2 × 10^−6^)^2^ ≈ 5.03 × 10^−11^*m*^2^.

Since the GUVs are filled with the same sucrose solution as the external medium, we need only consider the mass of the proteins and lipids in estimating the downward force due to gravity.

***We will take the Mass of one lipid to be around 800 Da, which translates into a mass of m_DOPC_***= 1.4 × 10^−24^ kg;

N_DOPC_ in 2 **μ**m radius GUV ∼ 10^7^ (<1 nm lipid)

Therefore, the total mass of lipid, ***M_lipid_*** = ***N_DOPC_*** x ***m_DOPC =_*** 10^-17^ kg

The mass of a single protein molecule with a molecular weight of 25 kDa is approximately 4.15 × 10^−23^ kg and it has an approximate radius of gyration = 2 *nm* = 2 × 10^−9^*m*

Since the proteins cover 2.5% of the Area of the GUV, **the number of proteins,** 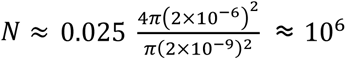 which is perhaps slightly overestimated since all the sites may not have proteins attached.

On the other hand, the bulk of the volume of the GUV is also filled with 50 µM protein solution. This gives a number contribution of

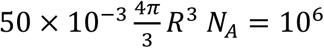

Thus, the **mass of all proteins** ≈ 8 × 10^−17^ kg. The **downward force due to gravity on the sGUV** is therefore, *W*_*np*_= 0.9 fN.

This weight of the sGUVs will lead to a sedimentation velocity of ***20*** nm***/s*** under Stokes conditions. On the other hand, the upward forces produced by enzyme activity can be estimated from Stokes law using the measured speeds (v =10 **μ**m/s):

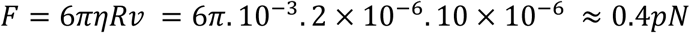

which is much larger than the fN weight. This implies that these sGUVs will not sink to the coverslip surface once the enzymes are activated. A similar value is obtained for the force if we use the measured *angular* speed instead.

#### **(b)** xGUVs (∼40 **µm** diameter)

The xGUVs on the other hand are ten times larger in radius than the sGUVs and hence the number of proteins on their surface is larger by a factor of 100. However, the number of proteins in the volume is larger by a factor 1000. This implies that their weight is

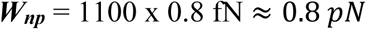

which leads to a sedimentation velocity of 2 **μ**m/s. Thus, we can expect them to quickly settle to the bottom.

In this case, the vertical forces due to enzyme activity must be determined from the measured angular velocity (*Ω* = 0.3/*s*) using Stokes equations (since these do not translate). This turns out to be

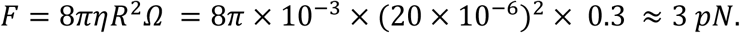

If we assume that forces of the same magnitude are applied in the horizontal direction, the velocity of the xGUV will be ≈8 **μ**m/s i.e., non-negligible. Thus, the reason for the absence of any translation must be sought elsewhere. We suspect that the estimation of force from the torque is incorrect for the reasons discussed by Ishikawa et al (41). Based on that work, we are led to include a factor of *ε* ∼ 10^-3^ which could potentially explain away the absence of translation.

So overall, these calculations are consistent with the observations that all xGUVs settle on the coverslip surface while the sGUVs are exuberantly dynamic.

### Tracking of the sGUVs

The trajectories of the GUVs were extracted using Particle Tracker 2D-3D of FIJI-ImageJ MosaicSuite and custom-made MATLAB code.

#### **(a)** Trajectory Analysis

To perform quantitative analysis of the trajectories, we evaluated the mean squared displacement (MSD), ⟨Δ***r***^**2**^(*t*)⟩ and velocity autocorrelation function (VACF), 〈***v***(*t*). ***v***(0)〉 for the sGUVs. The position coordinates of a GUV at a fixed time frame *t* is (x(*t*), y(*t*)). Then the MSD can be calculated as follows:

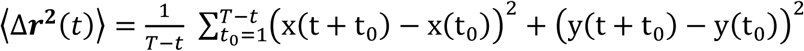

where *T* is the total number of time steps for a given sample trajectory.

For the VACF, the instantaneous velocities are given as: 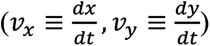. To calculate them, we discretize as follows: 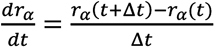, where *r_α_* (*t*) ≡ (*x*(t), y(t)) with *α* = (1,2). Δ*t* is the time step in our case, which is the inverse of the frame rate of our observations *i.e.* 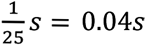. The VACF is given as:

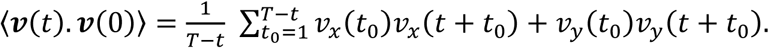

In **Figs. S18, S21 and S22** we plot VACF and MSDs for various sets of experiments and compare them with the corresponding theoretical expressions for a chiral active Brownian particle. By fitting these quantities with the cABP model, we extract the dynamical parameters such as self-propulsion velocity and the angular velocity of the GUVs.

#### (b) Dynamics of a chiral active GUV in two dimensions (2d)

In the special case, when the spinning axis of the GUV is perpendicular to the effective force, we get the following Langevin equations in two dimensions in the overdamped limit (*8,9,37,39*):

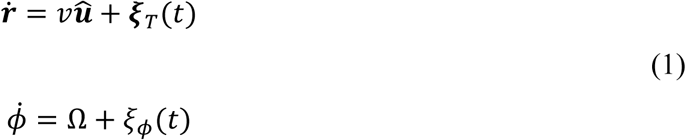

where, the position of the center of the GUV is r(t) and the point P makes an angle **ϕ**(t) at time t with a fixed axis **(Fig. S7)**. The dynamics is described using self-propulsion speed, 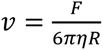 along the direction ***û*** of the point P and self-propelling angular speed, 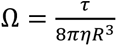. The translational noise *ξ*_*T*_ and the rotational noise *ξ*_*ϕ*_ have zero mean and variance given by, ⟨*ξ*_*Ti*_(*t*)*ξ*_*Tj*_(*t*^′^)⟩ = 2*Dδ*_*ij*_*δ*(*t* − *t*′) and, ⟨*ξ*_*ϕ*_(*t*)*ξ*_*ϕ*_(*t*^′^)⟩ = 2*D*_*r*_*δ*(*t* − *t*^′^). Here *D* and *D_r_* are the translational and rotational diffusion coefficients, respectively.

The various correlation functions of the cABP model can be determined exactly (*8,9,37,39*)

The orientation correlation takes the form:

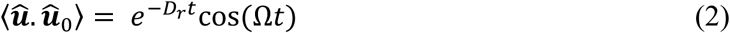

which shows decaying oscillations with a time scale *τ_r_* = 1*/D_r_* and a time period 2*π*/*Ω* set by the angular velocity (Ω) of the chiral GUV.

The velocity auto-correlation function can be also calculated to be:

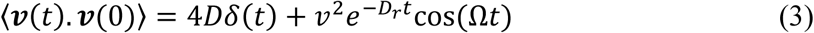

The mean squared displacement MSD(t) is obtained as:

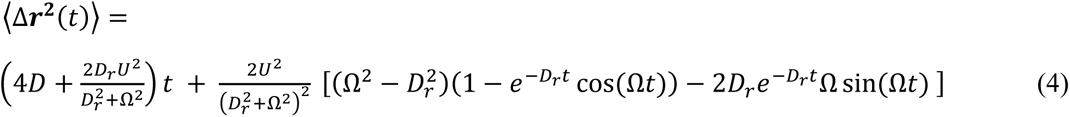

To understand the effect of chirality in MSD as observed in the experiments, we expand the MSD around *t* = 0 to get:

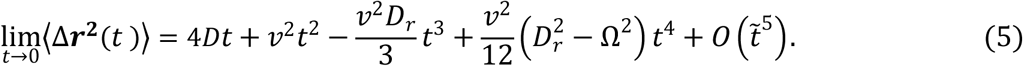

The effect of chirality is observed in the coefficient of *t*^4^ in this expression. This shows up as oscillations in the MSD (Fig. 5 in the main text & Figs. S18(b), S22).

We have followed the procedures mentioned below to evaluate the quantities associated with the chiral active motion. First, we obtain the frequency of the 〈***v***(*t*). ***v***(0)〉 by fitting with the sinusoidal function, *A*Sin(Ω*t* + *b*). Now, the time period of the correlation is 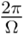, and in each period, the maximum value gives the values of the crests. Connecting these crests should obtain the curve, *v*^2^*e*^−*D_r_t*^ as we set 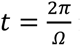 in eqn. (3). In cases such as ADK at low DSPC concentration, where no oscillations are observed in the MSDs, Ω is negligible, and 〈*v*(*t*). *v*(0)〉∼*v*^2^*e*^−*Drt*^ as expected for an ABP trajectory.

In **Fig. S22**, we plot the MSDs of the trajectories for which we have shown the velocity autocorrelation in **Fig. S21**. Using (3) we obtained the values of *v*,Ω, *D_r_*. D is then determined by fitting the MSDs, using the values of (*v*,Ω, *D_r_*) obtained previously. The resultant fits are shown in Figs. S18(b) and S22. There is good agreement between the experimental MSD and the theoretical expression (Eq. 4). It is worth mentioning that the MSDs show prominent oscillations in the intermediate time scales indicating the effect of chirality. This feature is not always observed in experiments of other cABPs.

---

**Fig. S1.**
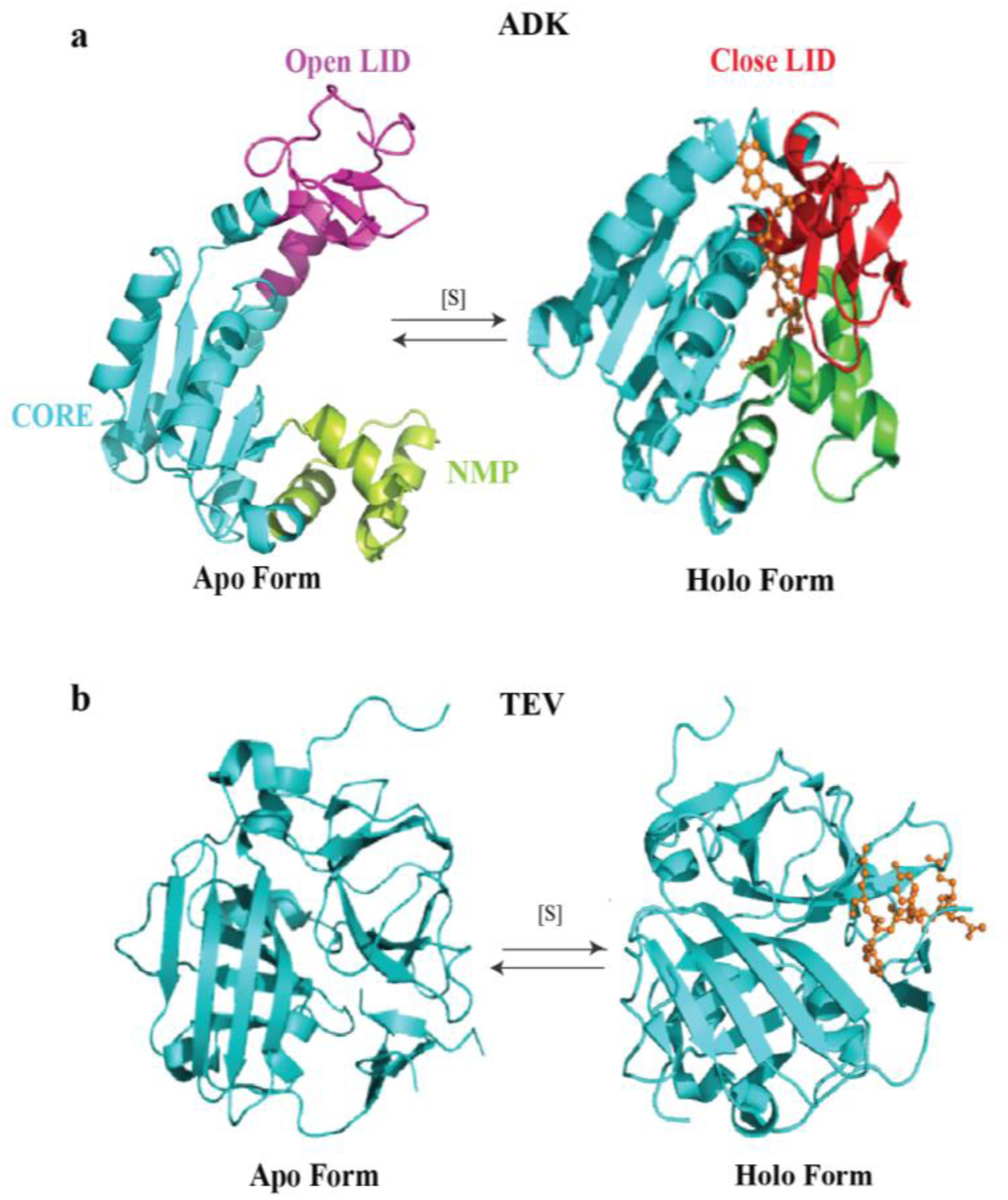
Conformational transitions in ADK and TEV. **(a)** Conformational transition of ADK from open LID (PDB ID: 4AKE, magenta) to closed LID (PDB ID: 1AKE, Red) upon substrate binding is presented using Ribbon diagram. **(b)** No change in conformation of TEV protease before (PDB ID: 1Q31) and after substrate binding (PDB ID: 1LVM) is illustrated using a ribbon diagram.

**Fig. S2.**
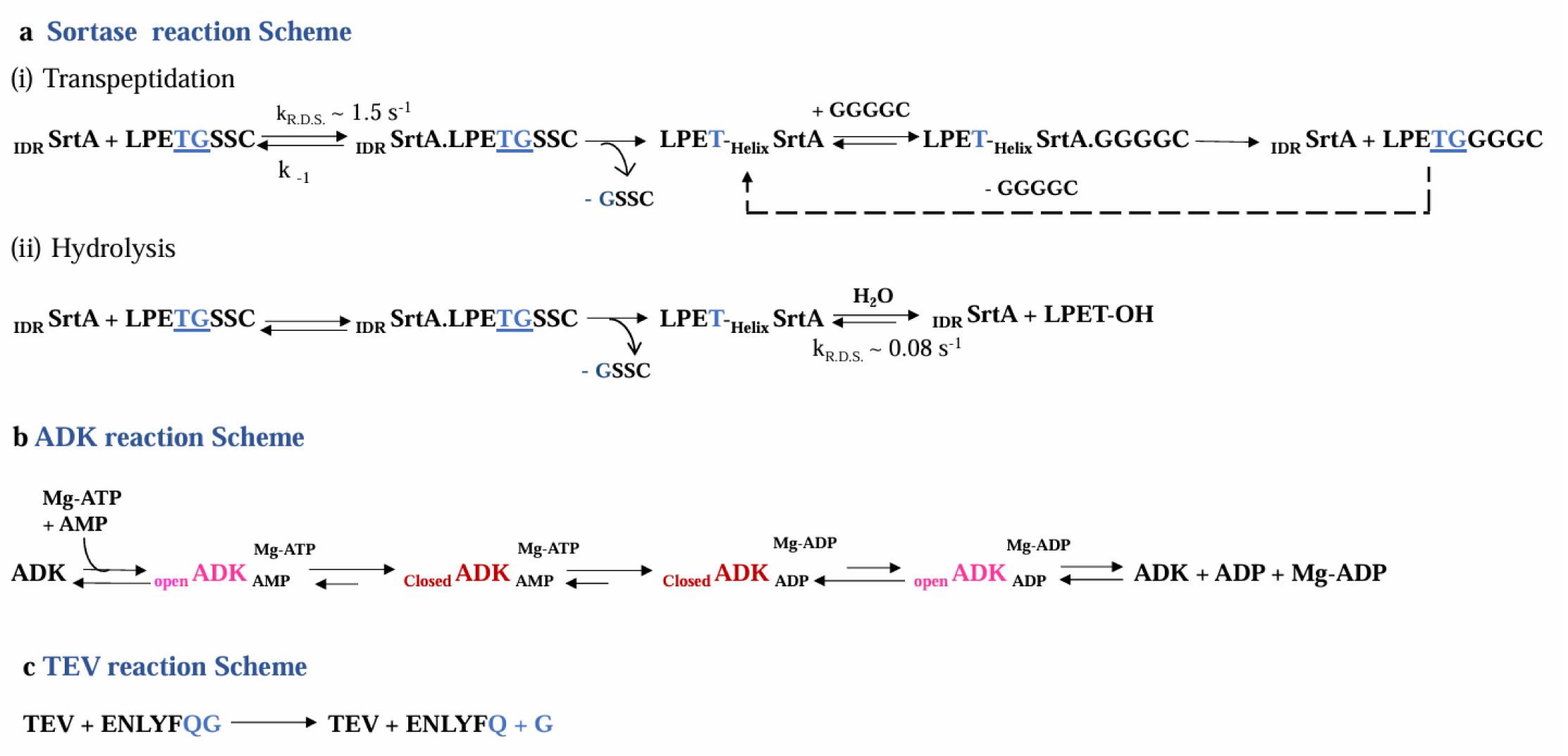
Reaction scheme for all three enzymes.

**Fig. S3.**
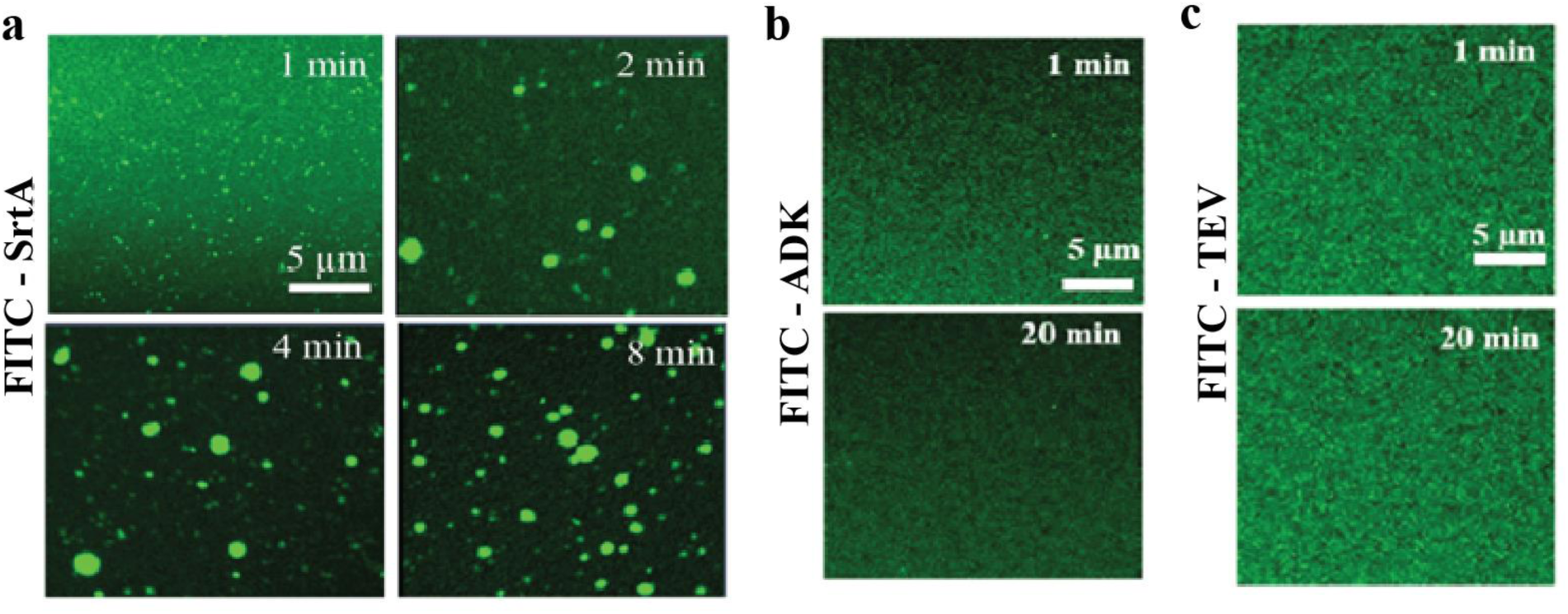
Spontaneous phase separation of SrtA in solution. **(a)** Time-lapse images of coalescing SrtA (15 µM) droplets. **(b-c)** Fluorescence images of ADK (30 µM) and TEV (30 µM) solutions respectively showed no phase-separation even after 20 mins. All images were recorded using a Confocal Fluorescence Microscope.

**Fig. S4.**
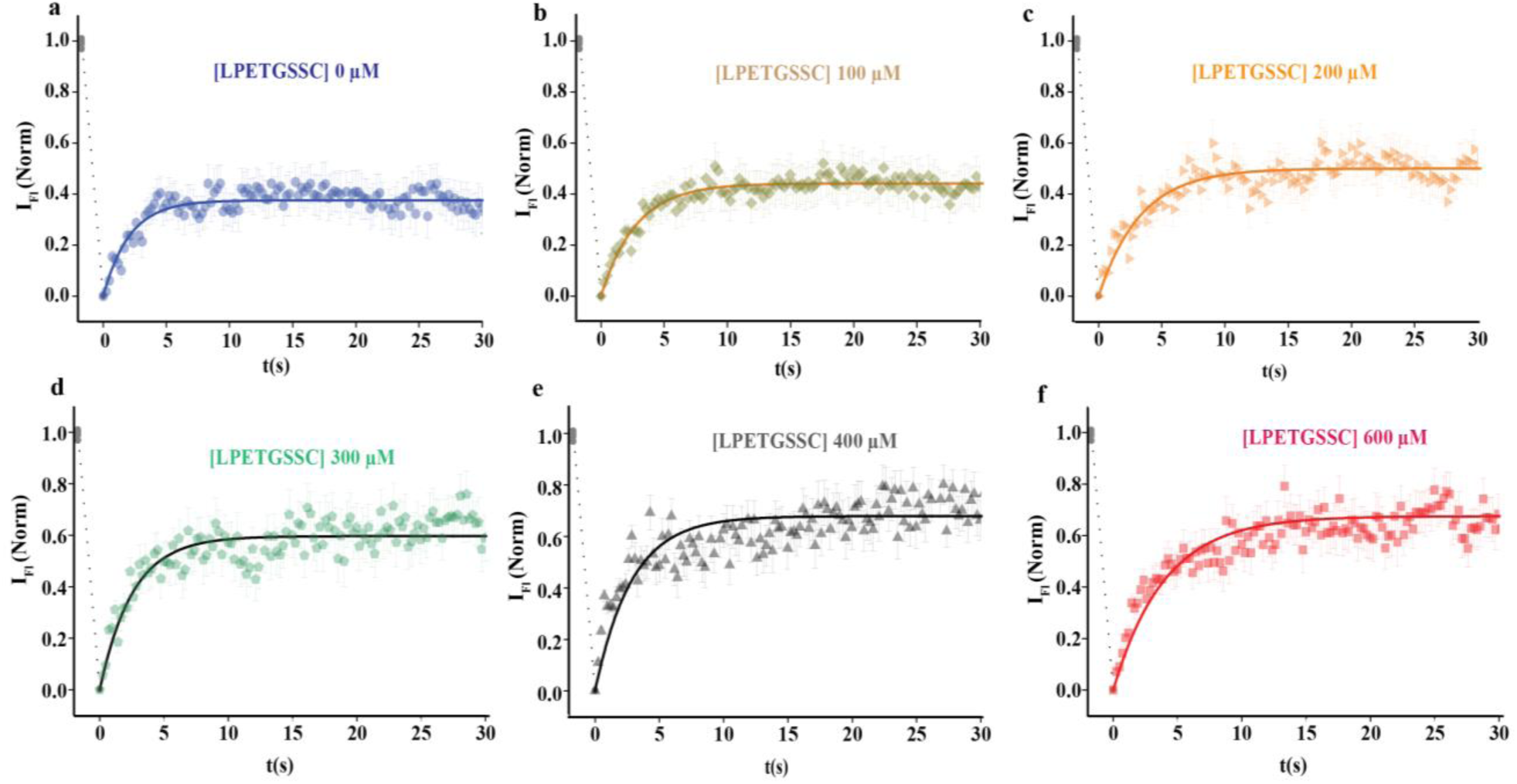
FRAP recovery data of SrtA within droplets at varying concentrations of - LPETGSSC-, **(a)** 0 µM, **(b)** 100 µM, **(c)** 200 µM, **(d)** 300 µM, **(e)** 400 µM, and **(f)** 600 µM are shown here. The solid lines represent the fitted curves. Three droplets (N=3) were analyzed for every condition.

**Fig. S5.**
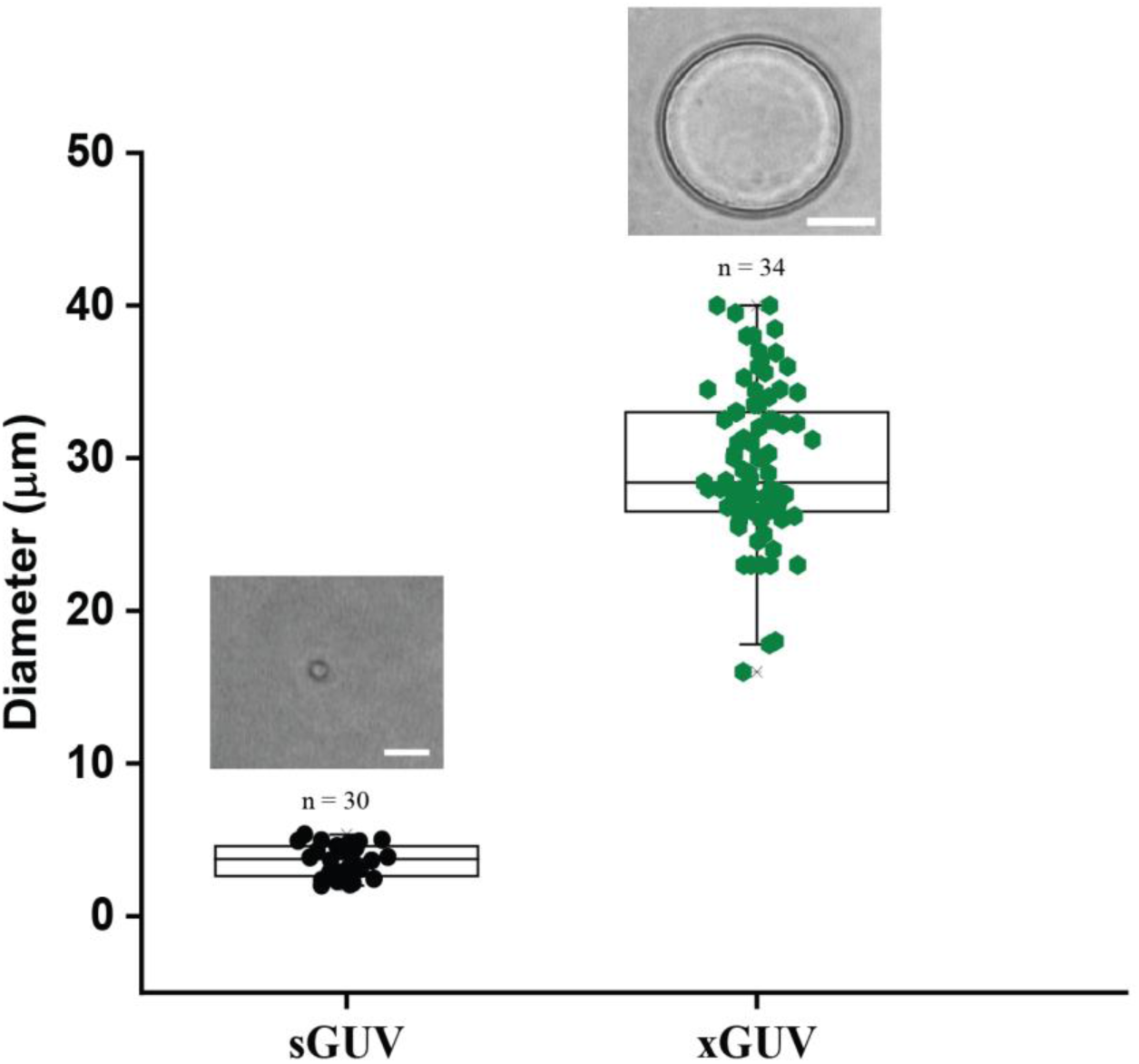
Size distribution of GUVs. This figure illustrates the size distribution of GUVs after they were eluted through a Sephadex G-50 column. The size estimates were derived from bright-field images of the GUVs, which are shown in the panels above each data point. Scale bar: 20 µm. Sample Size: ’n’ indicates the number of GUVs included in the analysis. Diameter of the GUVs was calculated using ImageJ software. The results indicate two main classes of GUVs: small GUVs (sGUVs) with average diameter of 3.4 ± 1.6 µm and large GUVs (xGUVs) with average diameter of 29 ± 6 µm.

**Fig. S6.**
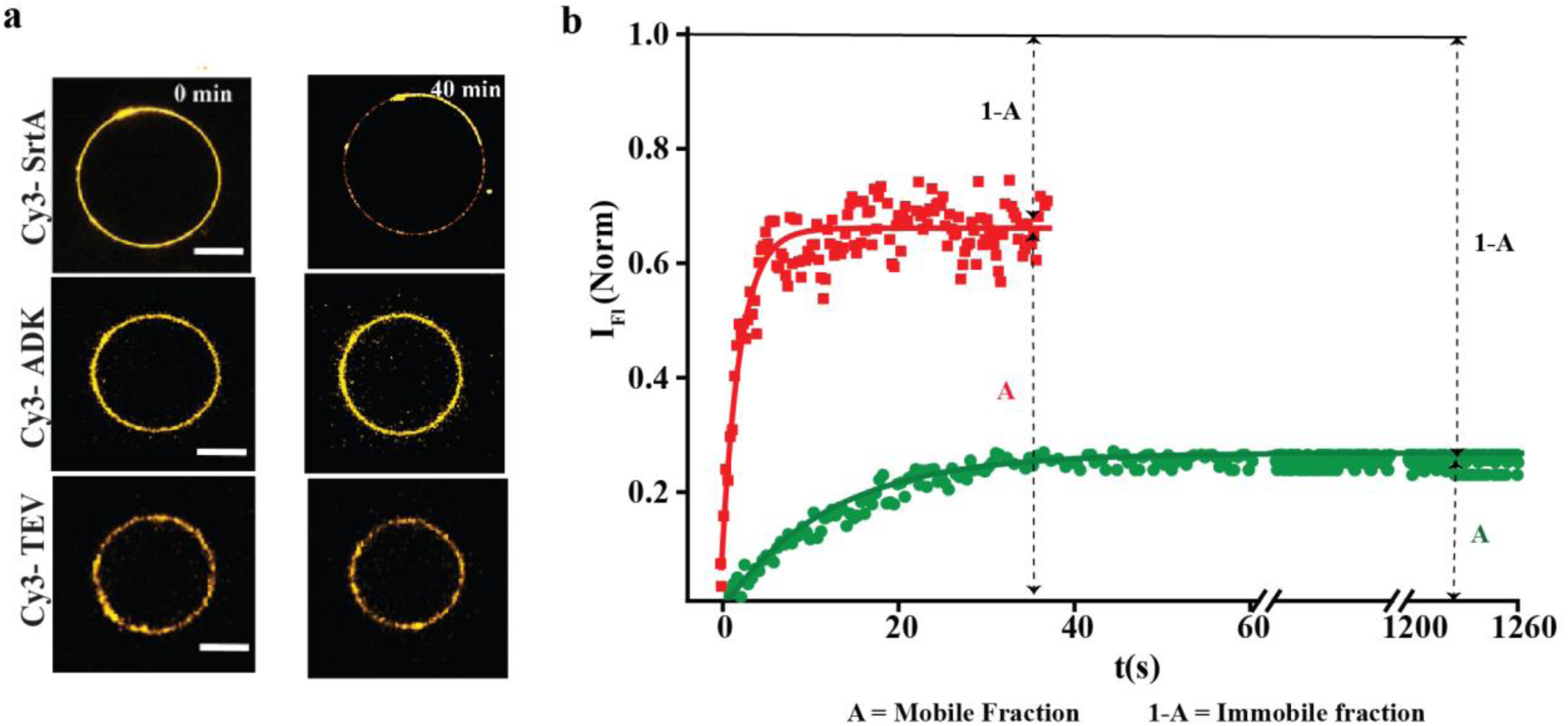
Enzyme-distribution on xGUVs. **(a)** The top panel displays fluorescence images taken with a confocal microscope, illustrating a gradual shift in SrtA distribution on the GUV membrane, leading to the formation of clusters. The middle and bottom panels show the distribution of ADK and TEV, respectively. Neither ADK nor TEV exhibits a clustering tendency and remains homogeneously distributed. Scale bars: 10 µm. (**b)** The temporal profile of FRAP for SrtA on GUVs is presented in green and TEV in red. The data indicate only a 20% recovery in intensity for SrtA even after 20 mins of recovery wait. The fluorescence intensity for TEV, in comparison, recovers nearly 70% within half minute, indicating that the lipid ordering by SrtA restricts lipid diffusion in GUVs.

**Fig. S7:**
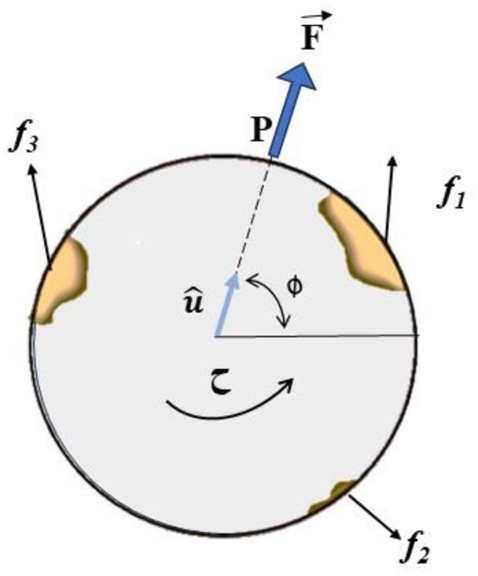
Force Diagram of a GUV. A schematic representation of the forces (*f_i_*) generated by the immobile clusters on a single GUV. F represents the effective radial force that acts at the fixed surface point P which makes an angle *ϕ* with an arbitrary axis. The torque *τ* is not necessarily in the plane of the figure.

**Fig. S8.**
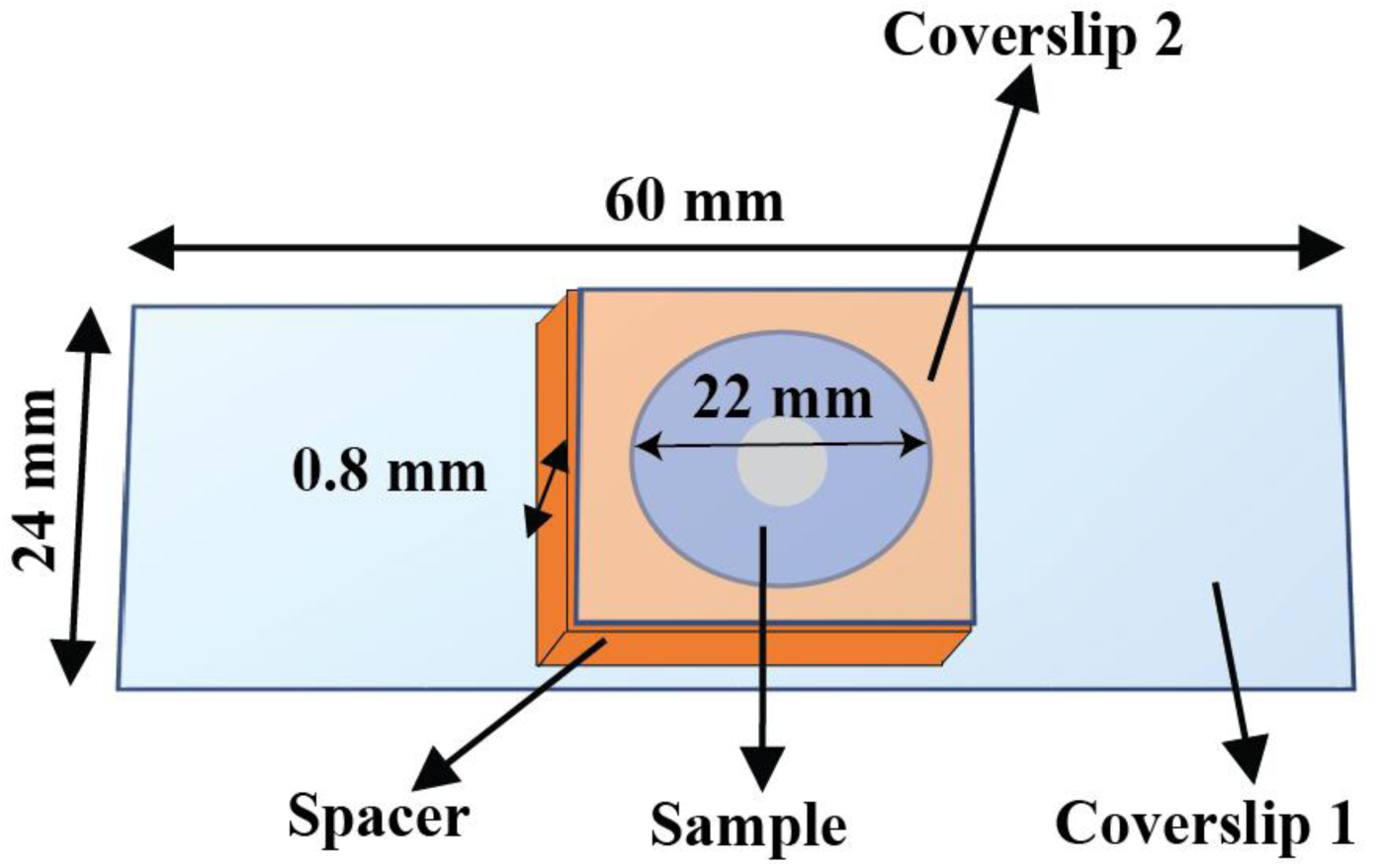
Pictorial description of the closed sample chamber used for inverted brightfield microscopy. All dimensions are indicated in the figure. The shaded region depicts the field of view (diameter 0.3 mm) recorded and analyzed.

**Fig. S9.**
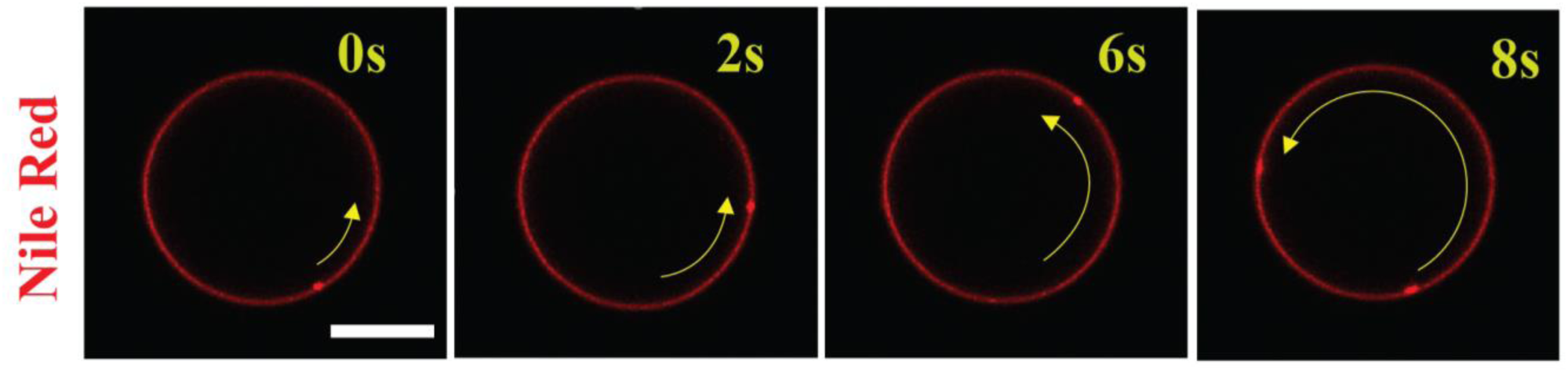
Activity of a SrtA coated GUV with catalytic turnover under an inverted fluorescence microscope. Time series snapshots of the movement of a patch of Nile red. Scale bar: 10 µm.

**Fig. S10.**
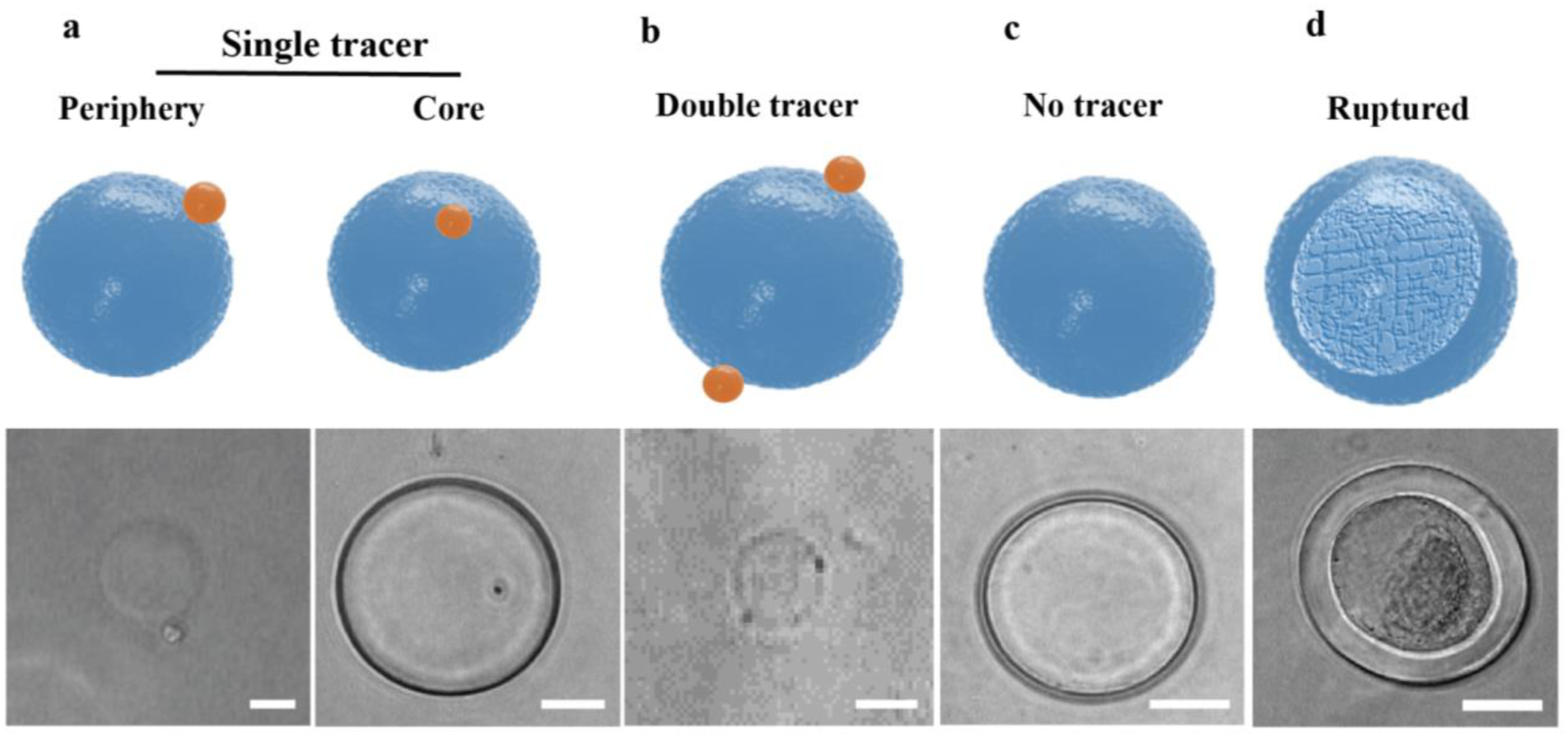
Non-specific attachment of passive tracers on GUV membrane. In the top panel, we pictorially describe the attachment of tracer on GUVs and the bottom panel reflects the corresponding Brightfield images. **(a)** Single tracer attached to the periphery (left) or the central region (right) of xGUVs. **(b)** Two tracers attached to GUVs membrane. **(c)** No tracer. **(d)** Representative image of a ruptured GUV. Scale bars: 10 µm.

**Fig. S11.**
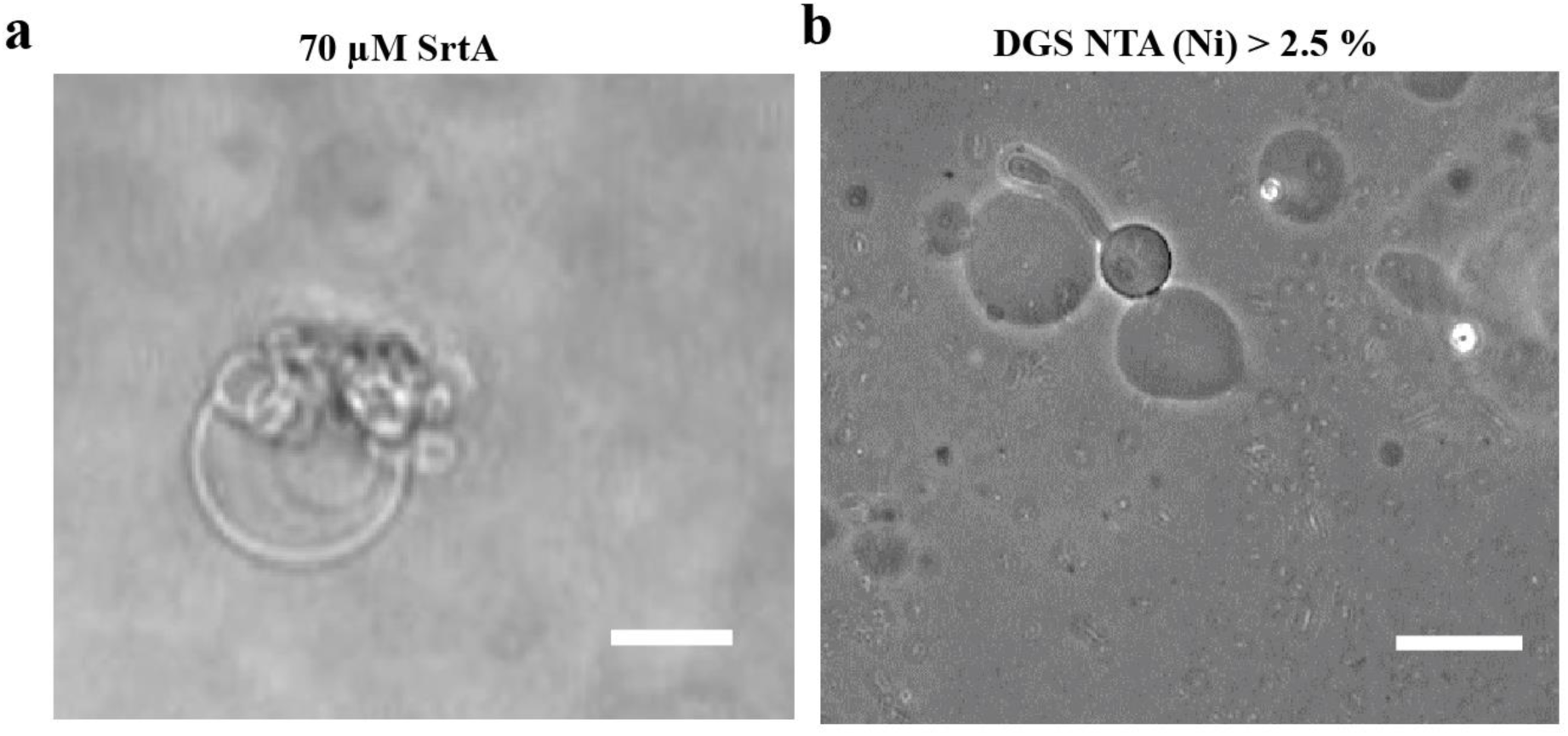
Distorted xGUVs. Typical images of deformed GUVs as seen for **(a)** increasing concentration of enzymes more than 50 µM on the membrane of GUVs, and with **(b)** increasing the percentage of DGS-NTA(Ni) beyond 2.5%. Scale bar: 20 µm.

**Fig. S12.**
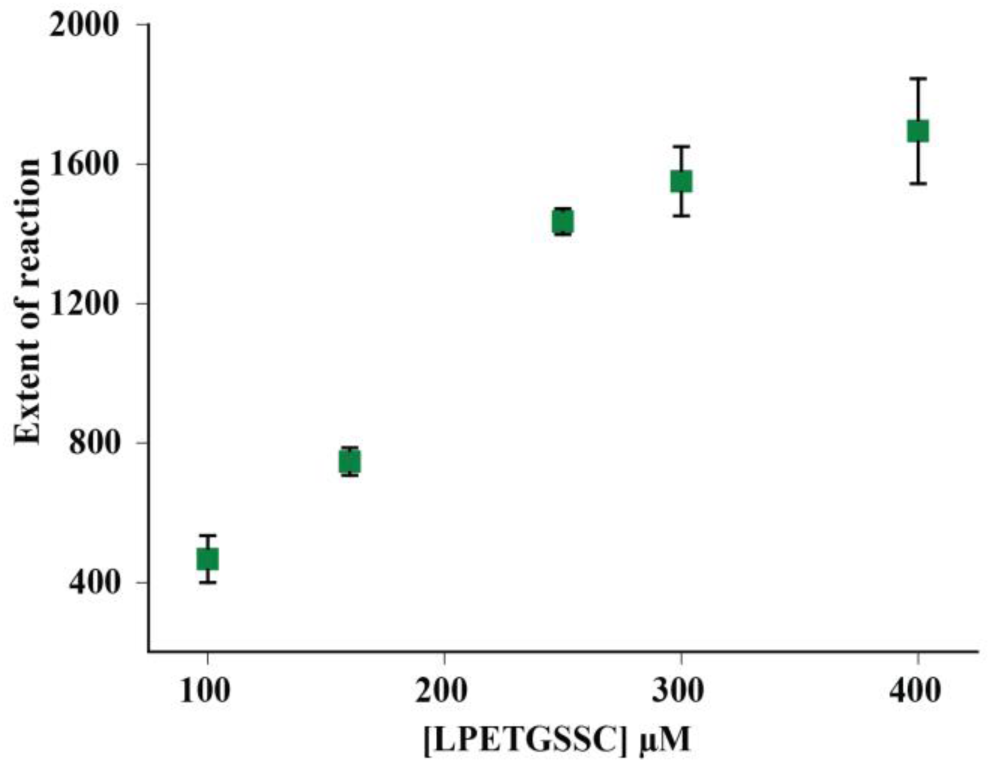
Variation in SrtA reactivity with signal peptide. Enzyme reactivity is calculated from the fluorescence intensity of MCA tagged -LPET- at t = 30 mins (**See Reaction Kinetics of SrtA in Solution with varying substrates in methods**). Same trend was observed when we measured the intensities at different time points of 20 min, 40 min, 60min.

**Fig. S13.**
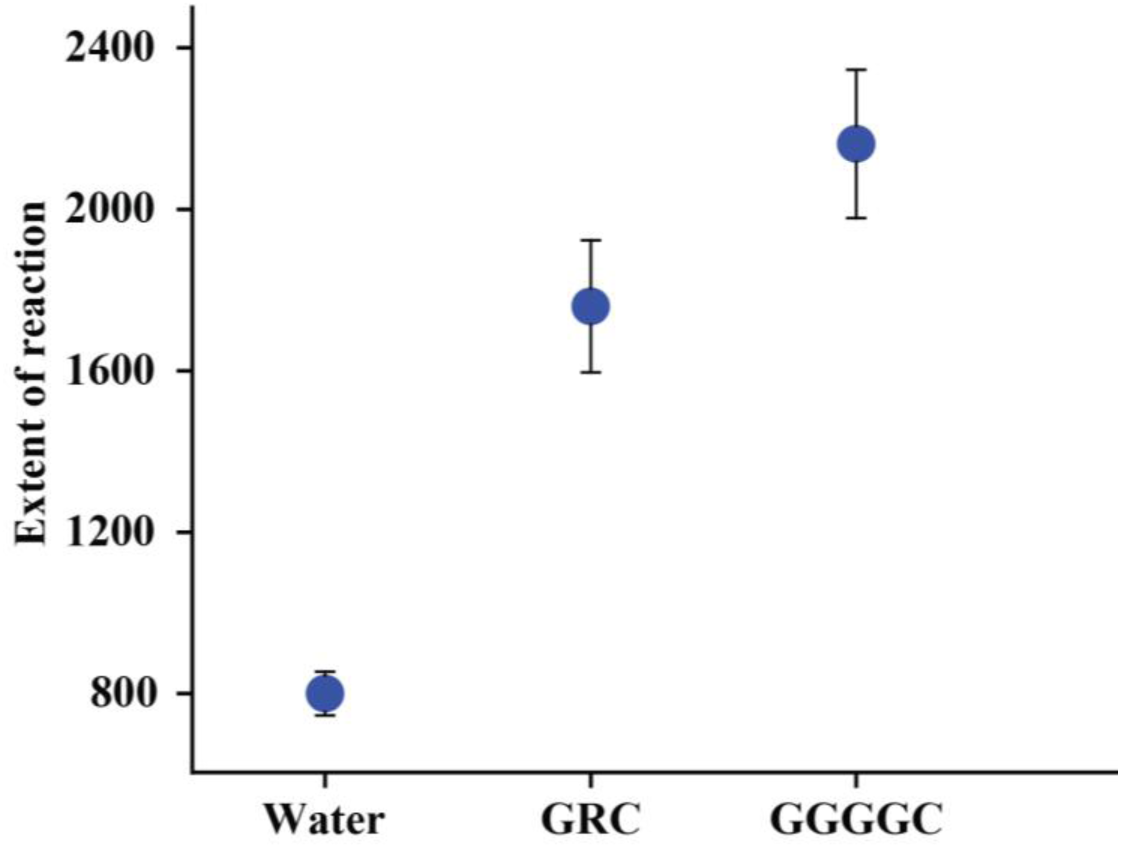
Variation in SrtA reactivity with nucleophiles. Enzyme reactivity is calculated from the fluorescence intensity of MCA tagged -LPET- at t = 30 mins with varying nucleophiles (**See Reaction Kinetics of SrtA in Solution with varying substrates in methods**). Same trend was observed when we measured the intensities at different time points (20 min, 40 min, 60min).

**Fig. S14.**
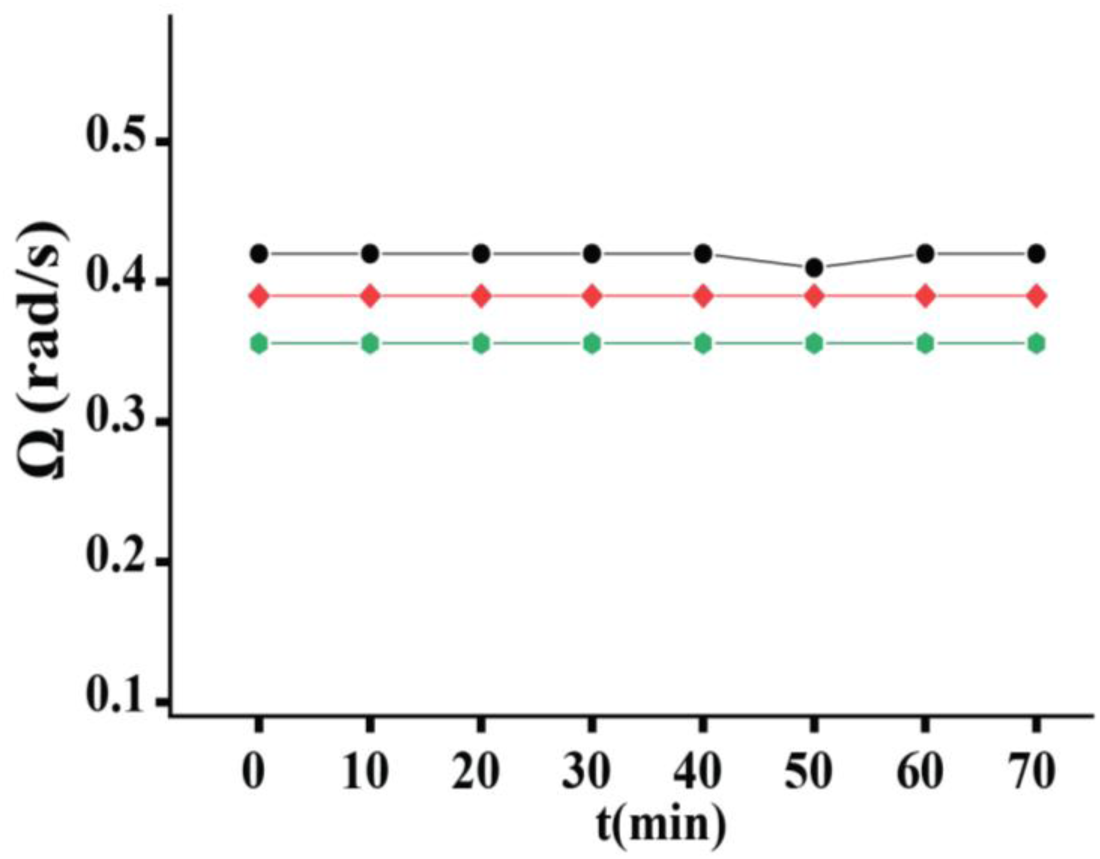
Constant rotation of xGUVs. No temporal change in the angular velocities of three different xGUVs are shown for 70 mins.

**Fig. S15.**
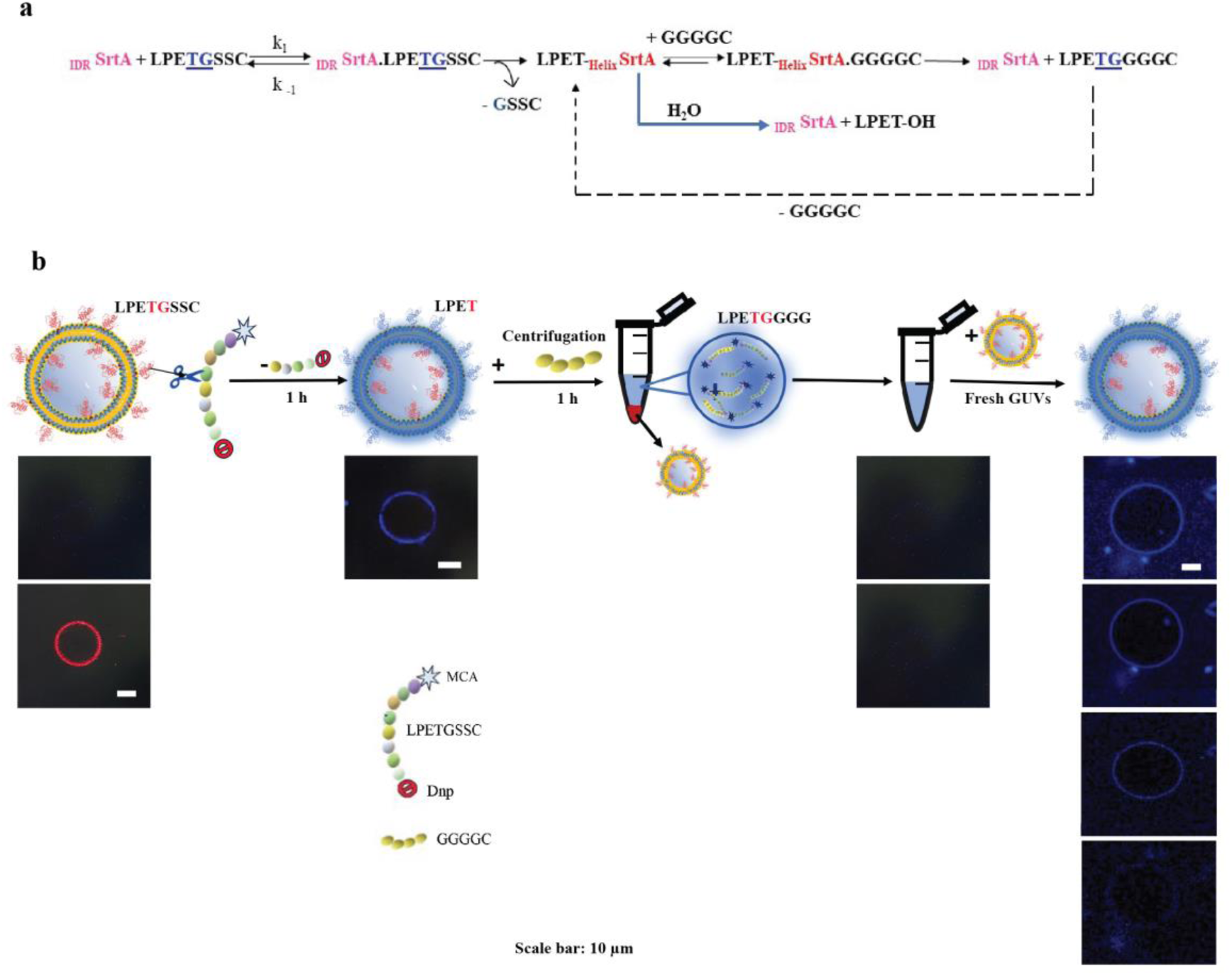
SrtA Utilizes Its Own Reaction Product as a Substrate. The top panel depicts the steps pictorially, while the bottom panel shows the corresponding outcomes in fluorescent images. To verify whether the final product from the SrtA-catalyzed turnover can serve as a substrate, we used the MCA-LPETGSSC-Dnp peptide, where MCA is a fluorophore attached at the N-terminus and Dnp is a quencher attached at the C-terminus. Without enzymatic cleavage, the signal peptide remains colorless, but upon excitation at 380 nm, it turns blue due to the cleavage of the T-G bond during thioesterification with SrtA. The experiment was carried out in the following steps:(i) SrtA-coated GUVs were incubated with 160 μM of MCA-LPETGSSC-Dnp (FRET-paired peptides).(ii) At 380 nm excitation, the GUVs appeared blue, indicating the cleavage of the T-G bond in the -LPETGSSC-peptide and the attachment of MCA-LPET to the SrtA coated GUVs.(iii) After centrifugation to remove unreacted peptides, a second substrate, GGGGC (360 μM), was introduced, and the blue color began to fade, suggesting the leaching of the product (MCA-LPETGGGGC) in solution.(iv) To remove the used GUVs, the sample was centrifuged for 30 minutes at 13,400 rpm. The supernatant was collected and examined under a microscope to confirm the absence of GUVs(v) Upon adding a fresh batch of SrtA-coated GUVs (50 μl) to the supernatant, the GUVs gradually turned blue, indicating that SrtA was using the product of the previous reaction (MCA-LPETGGGGC) as a substrate.

**Fig. S16.**
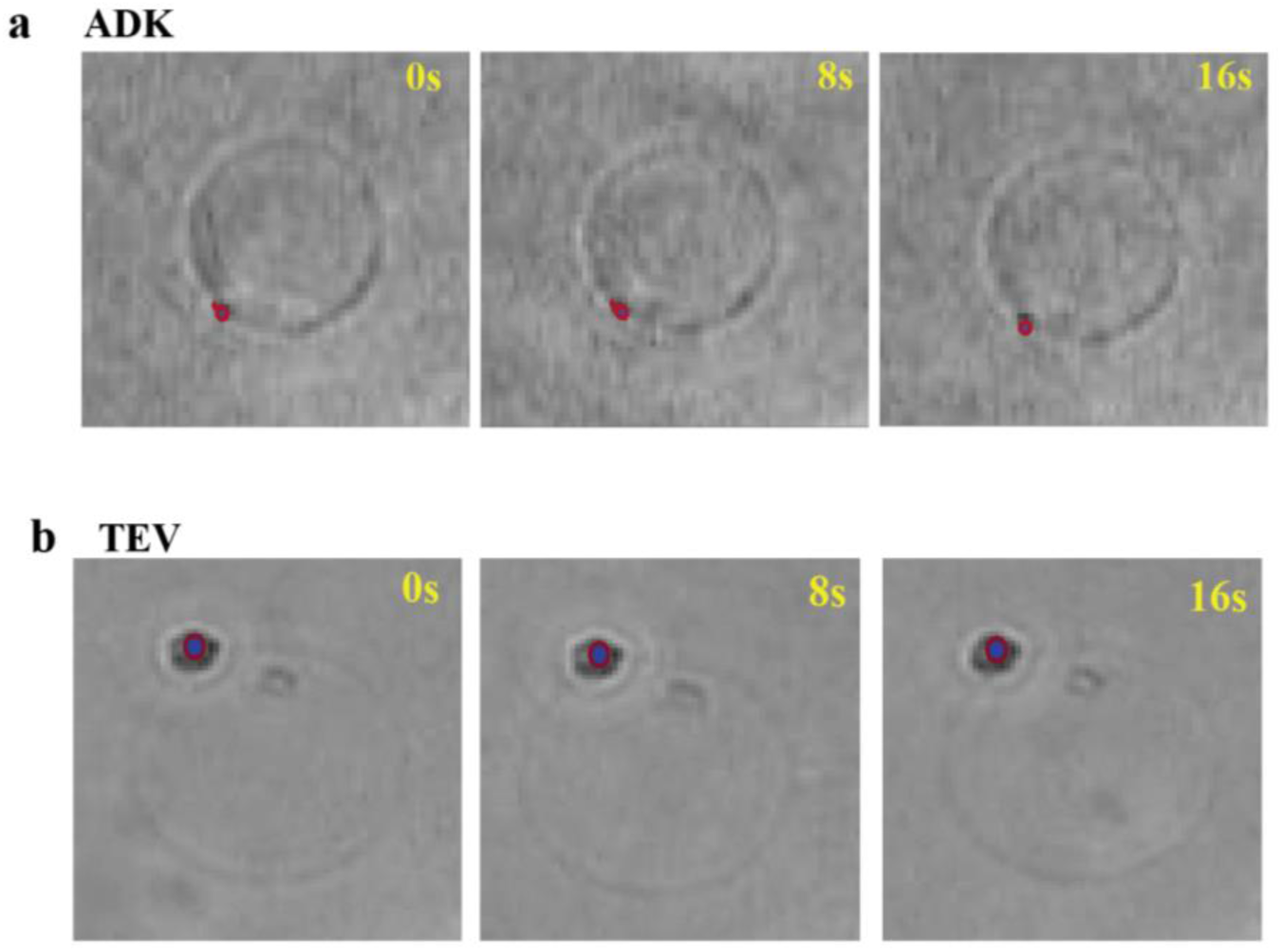
Passive xGUVs coated with ADK and TEV. No translational or rotational activity was noticed during catalysis in **(a)** ADK **(b)** TEV. Red and blue dots represent the position of tracers with time. Scale bar: 10 µm.

**Fig. S17.**
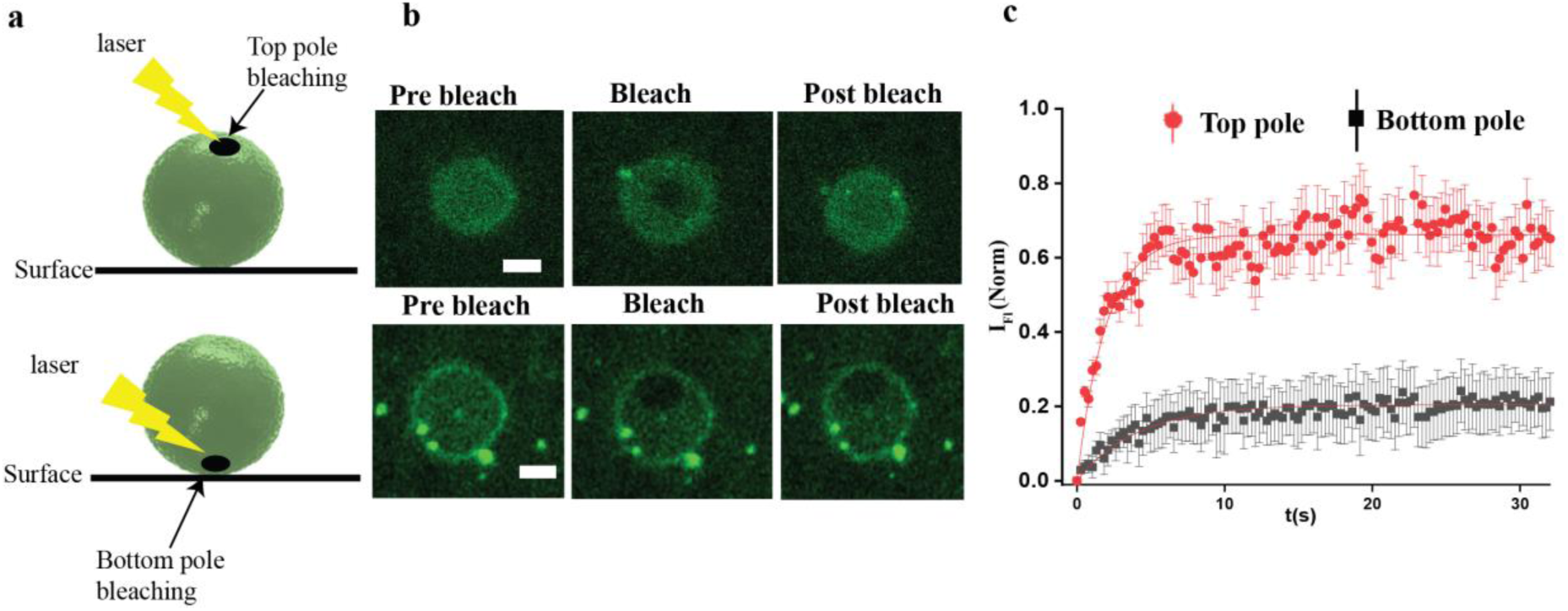
Lipid Diffusion at the Top and Bottom Pole of xGUVs. **(a)** Schematic illustration of the irradiated top and bottom pole of Giant Unilamellar Vesicles (GUVs). **(b)** Using a confocal microscope, we performed FRAP experiments on FITC-tagged TEV enzymes attached to the top pole (upper panel) and bottom pole (lower panel) of GUVs. The TEV enzymes were evenly distributed on the GUV surface. Representative FRAP images are shown. **(c)** The temporal profiles of fluorescence intensities after FRAP are plotted for the top pole (red) and bottom pole (black). The solid lines represent the fitted curves. For both cases, the analysis considered three GUVs (n = 3). Scale bar:10 µm.

**Fig. S18.**
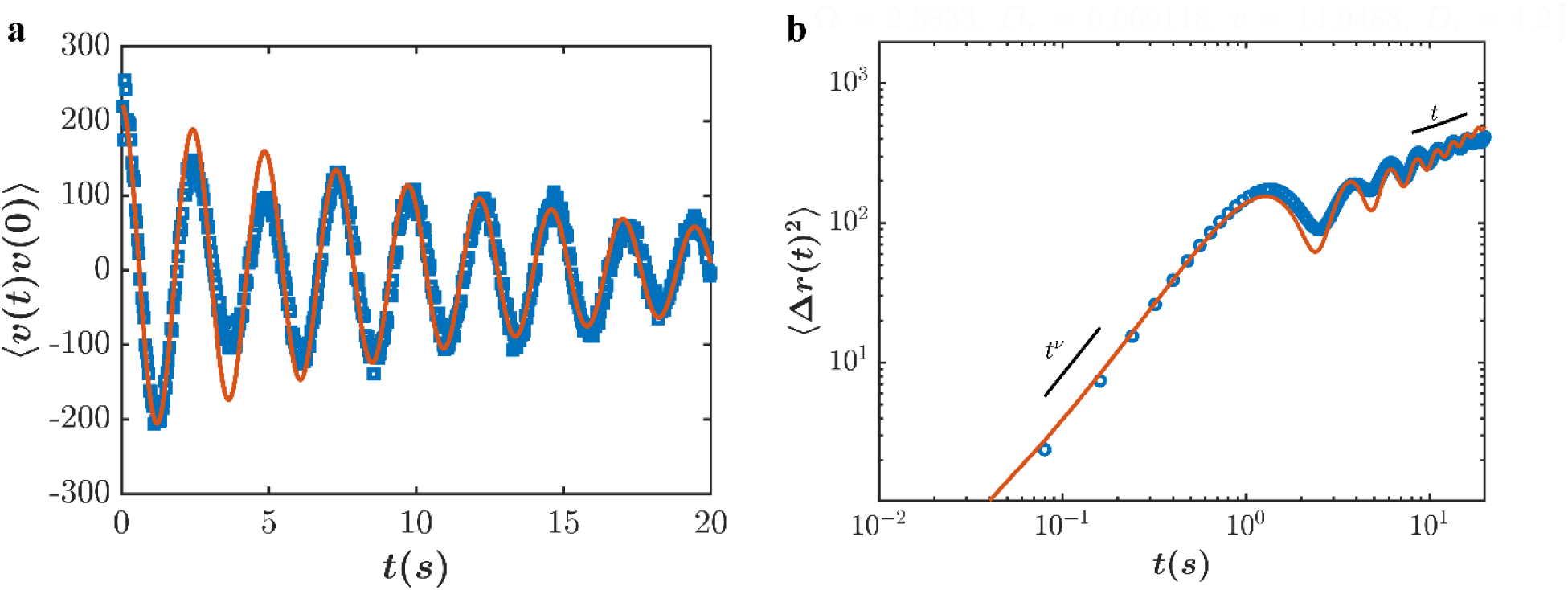
Velocity autocorrelation and MSD of SrtA-embedded sGUVs. **(a)** The velocity auto-correlation, ***v***(*t*)***v***(0) vs time, **t(s)** shows decaying oscillations. The ***v***(*t*)***v***(0) from the experiment in square dots shows good agreement with the cABP model shown in solid line, *v*^2^*e*^−*Drt*^cos (Ω*t*) (left). **(b)** We compare the MSD plots (right) from the experiment (in circles) with the MSD obtained from Eqn. (8) from the cABP model. MSD shows superdiffusive scaling in the initial regime *i.e. t* < 1*s* as MSD (*t*) ∼ *t*^*v*^ with *v* =1.65 (>1). In the long-time regime, *t* > 10*s* the MSD(*t*) is approaching diffusive scaling with MSD (*t*) ∼ *t*. The parameter values obtained from the fit are as follows: *v* = 14.9488 µ*m*/*s*, Ω = 2.5833 *rad*/*s*, *D*_*r*_ = 0.0691 *rad*^2^/*s*, *D* = 4.2 µ*m*^2^/*s*.

**Fig. S19.**
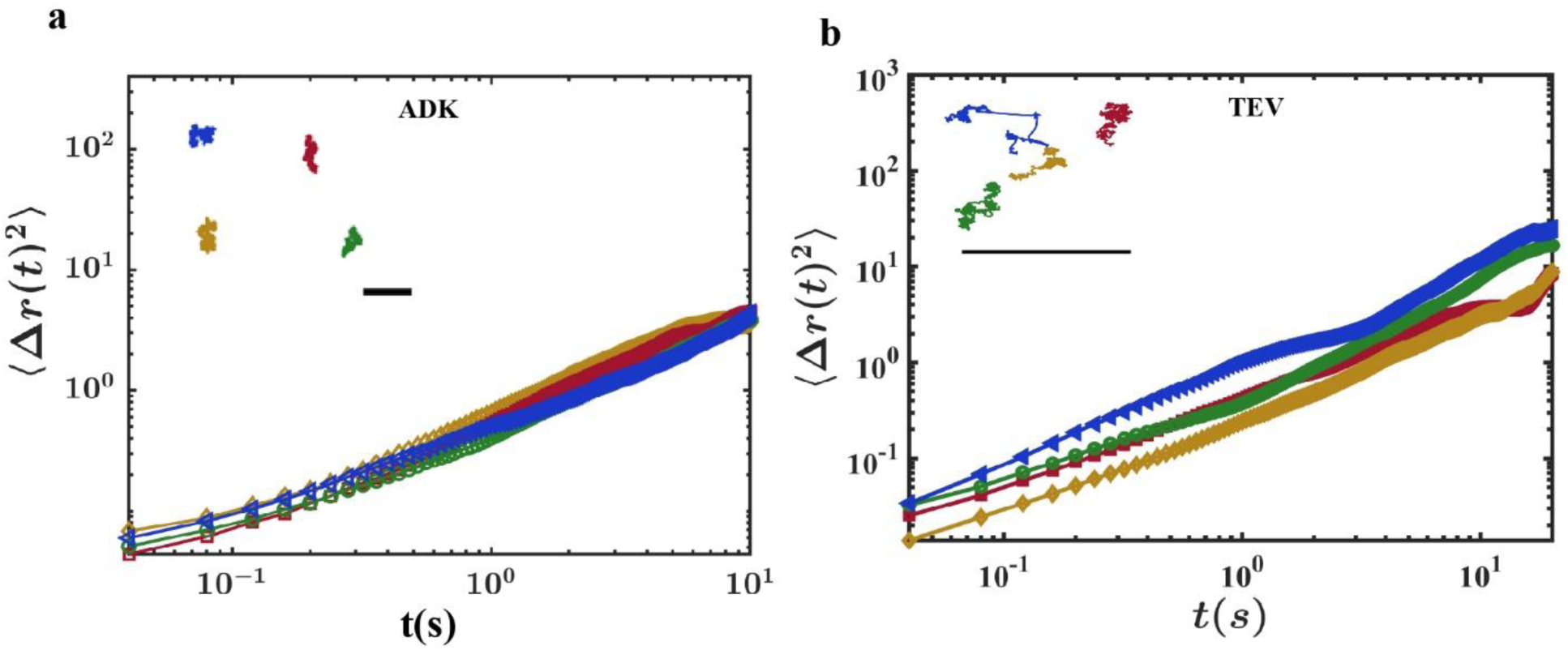
Passive motion of sGUVs. **(a)** MSD plot of ADK embedded sGUVs. **(b)** MSD plot of TEV embedded sGUVs. Insets show the corresponding trajectories of the sGUVs. Scale bar: 10 µm. The MSDs show purely diffusive features indicating the lack of activity.

**Fig. S20.**
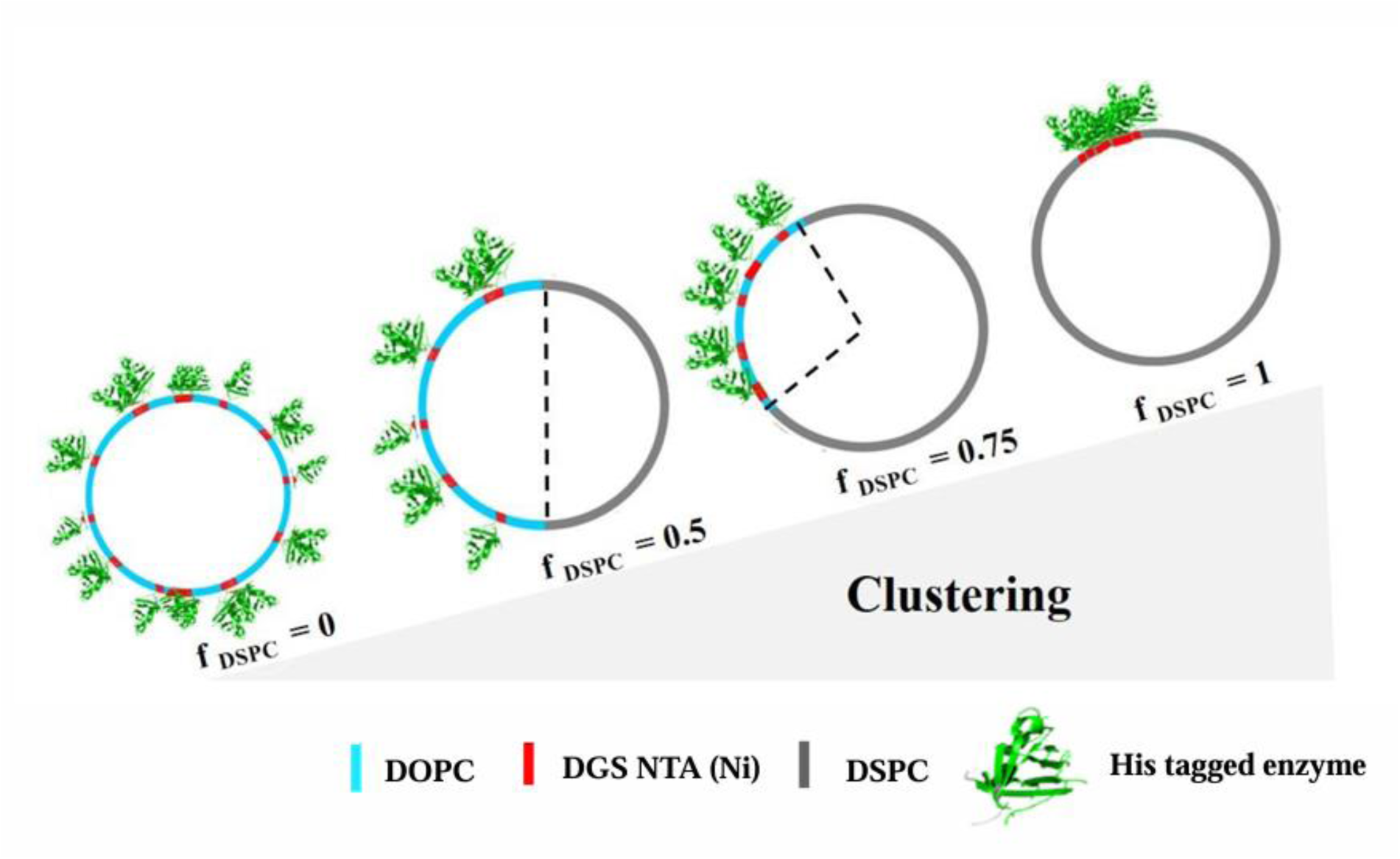
DSPC induced clustering on the GUVs: Sketch represents the pictorial distribution of enzymes on the surface of GUVs with increasing f_DSPC_.

**Fig. S21.**
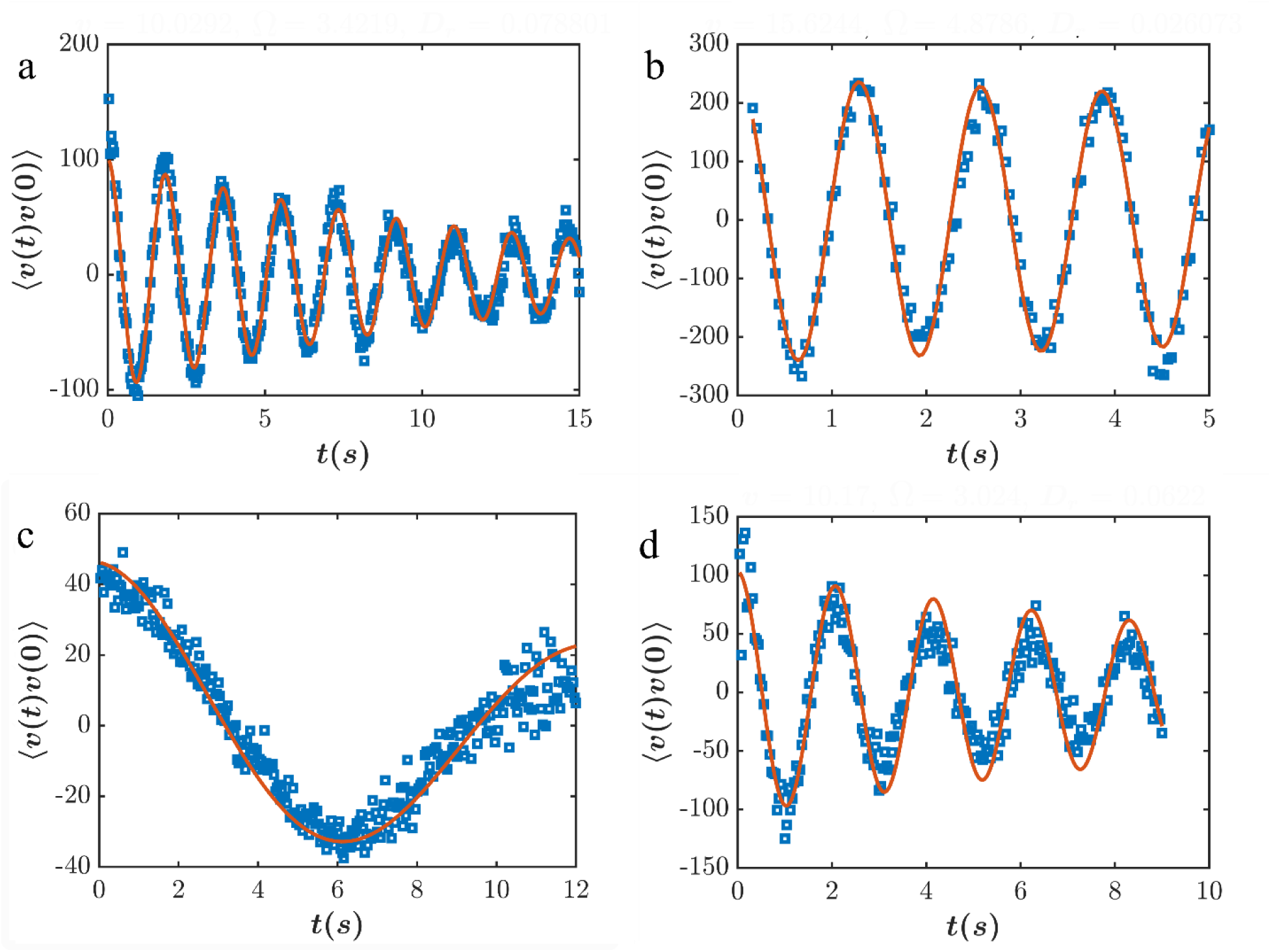
Sample plots of velocity correlation with time for different enzymes with different DSPC concentrations. (a -. **b)** show the for sortase at *f_DSPC_* = 0.5 and 1 and **(c-d)** represent the velocity correlation of ADK at *f_DSPC_* = 0.5 and 1 respectively. We compare Eq. (3) with the velocity correlation plots to extract the self-propulsion velocity, *v* and the angular velocity, Ω along with the rotational diffusion constant, *D*_*r*_. Using the extracted quantities, we show in Fig.S22 the comparison of MSDs obtained from the experiments with the MSD Eqn. (4) from the cABP model.

**Fig. S22.**
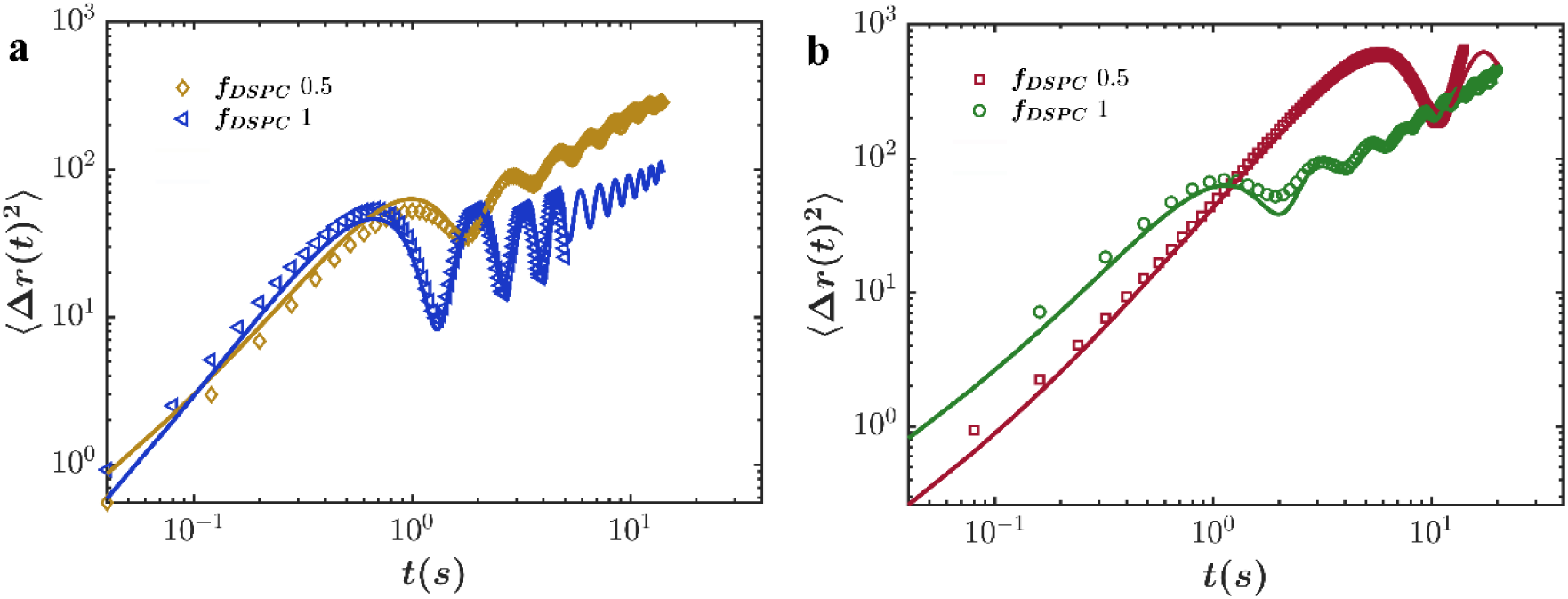
Mean squared displacement, corresponding to the velocity autocorrelation shown in **Fig.S21**. The points represent the MSD of SrtA (a) and ADK(b) obtained from the trajectory of sGUV in the experiment. The solid line represents the expression in Eq. (4), where *v*, Ω and D_r_ *a*re obtained from the fit shown in **Fig**.**S21**. *D* is obtained by a fit to the MSD. (**a**) *f_DSPC_* = 0.5, *v* = 11.85 *μm*/*s*, Ω = 3.43*rad*/*s*, *D*_*r*_ = 0.08*rad*^2^/*s*, *D* = 4.1 *μm*^2^/*s*, (**a**)*f_DSPC_* = 1.0, *v* = 15.79 *μm*/*s*, Ω = 4.8 *rad*/*s*, *D*_*r*_ = 0.03 *rad*^2^/*s*, *D* = 1.24 *μm*^2^/*s* (**b**)*f_DSPC_* = 0.5, *v* = 6.8 *μm*/*s*, Ω = 0.50 *rad*/*s*, *D*_*r*_ = 0.06 *rad*^2^/*s*, *D* = 1.23 *μm*^2^/*s* (**b**)*f_DSPC_* = 1.0, *v* = 10.17*μm*/*s*, Ω = 3.02 *rad*/*s*, *D*_*r*_ = 0.06 *rad*^2^/ *s*, *D* = 4.05*μm*^2^/*s*.

**Fig. S23.**
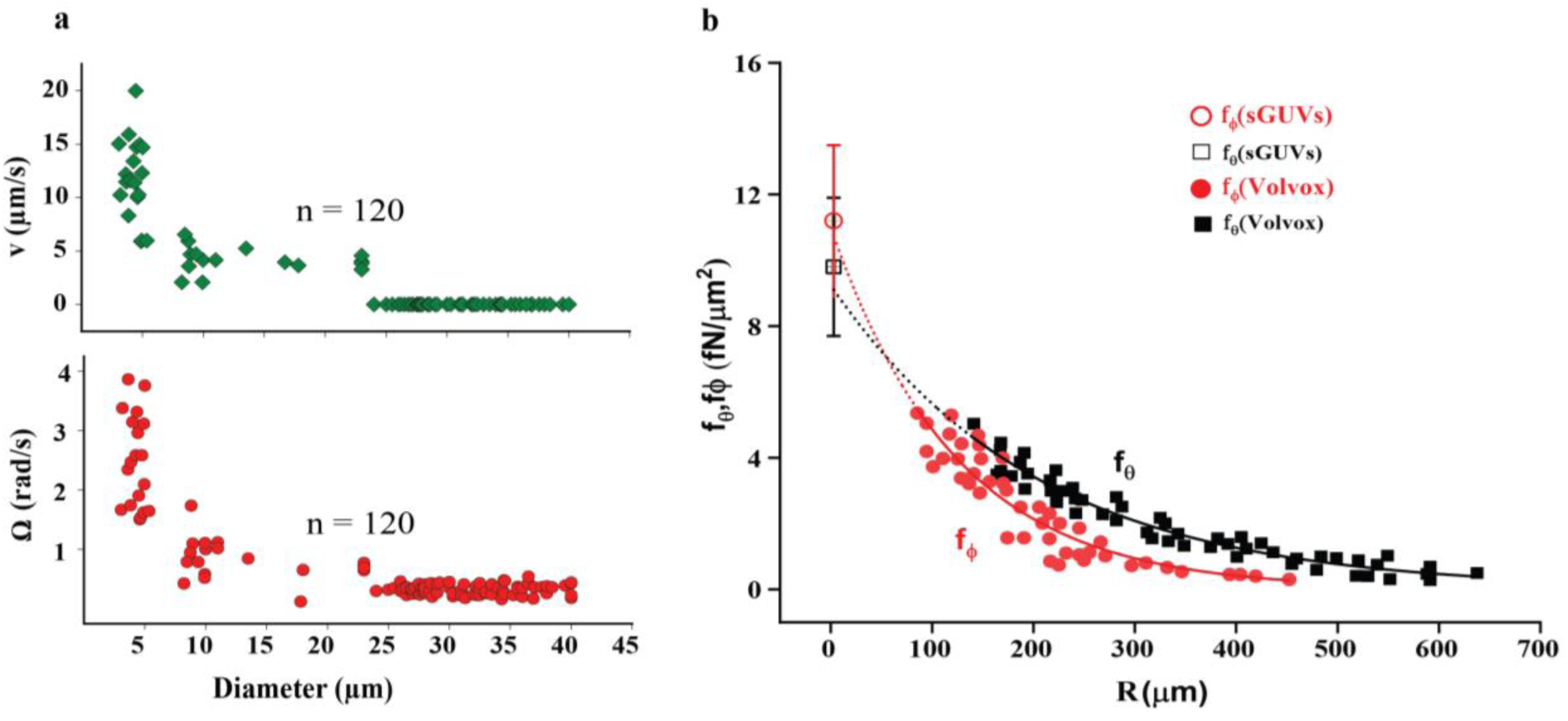
Variations in the dynamics of **μ**-rotors with size and the comparison of shear stresses generated by the gyrating sGUV with Volvox. **(a)** The linear speed (v) and angular velocity (Ω) drops with increasing GUV diameter. **(b)** Plot of shear stresses generated by Volvox (100 - 500 microns) and sGUVs (1-5 microns). Volvox generates a uniform tangential shear stress on the sphere’s surface through periodic and non-reciprocal flagellar beating, with shear stress components defined as, 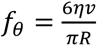 (where R is radius, η is viscosity, and v is the linear velocity) acting in the direction of the polar angle θ and 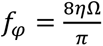 acting in the direction of the azimuthal angle ϕ. Data is sourced from **Reference** (**3**) with permission. The shear stresses components, *f*_*θ*_ & *f*_*φ*_, for Volvox are fitted to exponential decay and extrapolated (dotted line) to a radius of 3 µm. A striking resemblance in the shear stresses generated by our floating µ-rotors and giant Volvox is observed.

